# Herpes simplex virus-1 pUL56 degrades GOPC to alter the plasma membrane proteome

**DOI:** 10.1101/729343

**Authors:** Timothy K. Soh, Colin T. R. Davies, Julia Muenzner, Viv Connor, Clément R. Bouton, Henry G. Barrow, Cameron Smith, Edward Emmott, Robin Antrobus, Stephen C. Graham, Michael P. Weekes, Colin M. Crump

**Affiliations:** Division of Virology, Department of Pathology, Cambridge University, Cambridge, CB2 1QP, UK; Cambridge Institute for Medical Research, Cambridge University, Cambridge, CB2 0XY, UK

**Keywords:** Herpesvirus, Virus host interaction, Immune evasion, Membrane trafficking, Proteasomal degradation, Quantitative proteomics, Uncharacterized ORF

## Abstract

Herpesviruses are ubiquitous in the human population and they extensively remodel the cellular environment during infection. Multiplexed quantitative proteomic analysis over a whole time-course of herpes simplex virus (HSV)-1 infection was used to characterize changes in the host-cell proteome and to probe the kinetics of viral protein production. Several host-cell proteins were targeted for rapid degradation by HSV-1, including the cellular trafficking factor GOPC. We identify that the poorly-characterized HSV-1 protein pUL56 binds directly to GOPC, stimulating its ubiquitination and proteasomal degradation. Plasma membrane profiling revealed that pUL56 mediates specific changes to the surface proteome of infected cells, including loss of IL18 receptor and Toll-like receptor 2, and delivery of Toll-like receptor 2 to the cell-surface requires GOPC. Our study highlights an unanticipated and efficient mechanism whereby a single virus protein targets a cellular trafficking factor to modify the abundance of multiple signaling molecules at the surface of infected cells.

## Introduction

Herpesviruses are ubiquitous in the human population and are characterized by an ability to establish lifelong infections. Greater than two thirds of the world population are estimated to be infected with HSV-1 and HSV-2 (Looker et al., 2008; Looker et al., 2015). These infections are generally asymptomatic or give rise to mild symptoms following viral reactivation (oral or genital sores), although they can cause severe diseases of the eye (herpes keratitis), central nervous system (herpes encephalitis), or systemic infections in those with compromised or immature immune systems (Gnann and Whitley, 2017; Koujah et al., 2019; Pinninti and Kimberlin, 2018).

The replication cycle of herpesviruses entails a complex and carefully controlled transcriptional cascade of viral genes that function both to generate infectious particles and to modulate host factors. HSV-1 genes are conventionally separated into three broad temporal classes (immediate-early, early, and late), where proteins expressed earliest during infection serve as transcription factors and/or modulate the host-cell environment and immune responses, while those expressed late are structural components of the virion. The best-studied HSV-1 immunomodulatory proteins are infected cell protein 0 (ICP0) and virion host shutoff protein (vhs). These proteins are known to modulate the host-cell proteome by suppressing the expression and/or promoting the degradation of various host proteins (Boutell et al., 2011; Chelbi-Alix and de The, 1999; Jiang et al., 2016; Lees-Miller et al., 1996; Lilley et al., 2011; Orzalli et al., 2013; Su and Zheng, 2017; Zenner et al., 2013). However, the global temporal effects of HSV-1 replication on the host proteome remain poorly characterized. To date there has been one large-scale proteomic analysis of HSV-1 infection. This work, performed in fibroblasts, quantified the abundance of approximately 4000 host proteins and characterized changes in protein post-translational modification following infection (Kulej et al., 2017). However, the molecular mechanisms underlying these changes were not characterized.

We developed quantitative temporal viromics (QTV) as a method to enable highly-multiplexed quantitative analysis of temporal changes in host and viral proteins throughout the course of a productive infection (Weekes et al., 2014). QTV employs tandem mass tags (TMT) and MS3 mass spectrometry to facilitate precise quantitation of each protein, and we have applied this technique to study several viruses including human cytomegalovirus (HCMV), Epstein-Barr virus, vaccinia virus, and BK polyomavirus (Caller et al., 2019; Ersing et al., 2017; Soday et al., 2019; Weekes et al., 2014).

We have now performed QTV analysis throughout a single replication cycle of HSV-1 in human keratinocytes, the natural target of HSV-1 lytic infection. At each time point we quantified almost 7000 human proteins and >90% of canonical HSV-1 proteins, and we have found evidence for protein expression from 17 novel HSV-1 open reading frames (ORFs). We have identified host proteins that are rapidly degraded by HSV-1, including the cellular trafficking factor Golgi associated PDZ and Coiled-coil motif containing protein (GOPC). Further, we demonstrate that GOPC degradation is mediated by the poorly-characterized HSV-1 protein pUL56. Plasma membrane profiling and flow cytometry show that pUL56-mediated degradation of GOPC reduces the cell-surface abundance of multiple host proteins, including the immune signaling molecule Toll-like receptor 2 (TLR2). This highlights an unanticipated and highly-efficient mechanism whereby HSV-1 specifically targets a cellular trafficking factor in order to manipulate the abundance of multiple proteins on the surface of infected cells.

## Results

### Quantitative Temporal Viromic study of HSV-1 infection

To construct an unbiased global picture of changes in host and viral proteins throughout the course of HSV-1 infection, we infected human keratinocyte cells (HaCaT) with HSV-1 at a high multiplicity of infection (10 PFU/cell) (Figure 1, Table S1). Immunofluorescence analysis of parallel samples confirmed that >95% of cells were infected. Ten-plex TMT and MS3 mass spectrometry were used to quantify changes in protein expression over seven time points (Figure 1A). A particular advantage of such TMT-based quantitation is the measurement of each protein at every time point. This generated the most complete proteomic dataset examining the lytic replication cycle of HSV-1 to date, quantifying 6956 human proteins and 67/74 canonical HSV-1 proteins, and provided a global view of changes in protein expression during infection.

**Figure 1:**
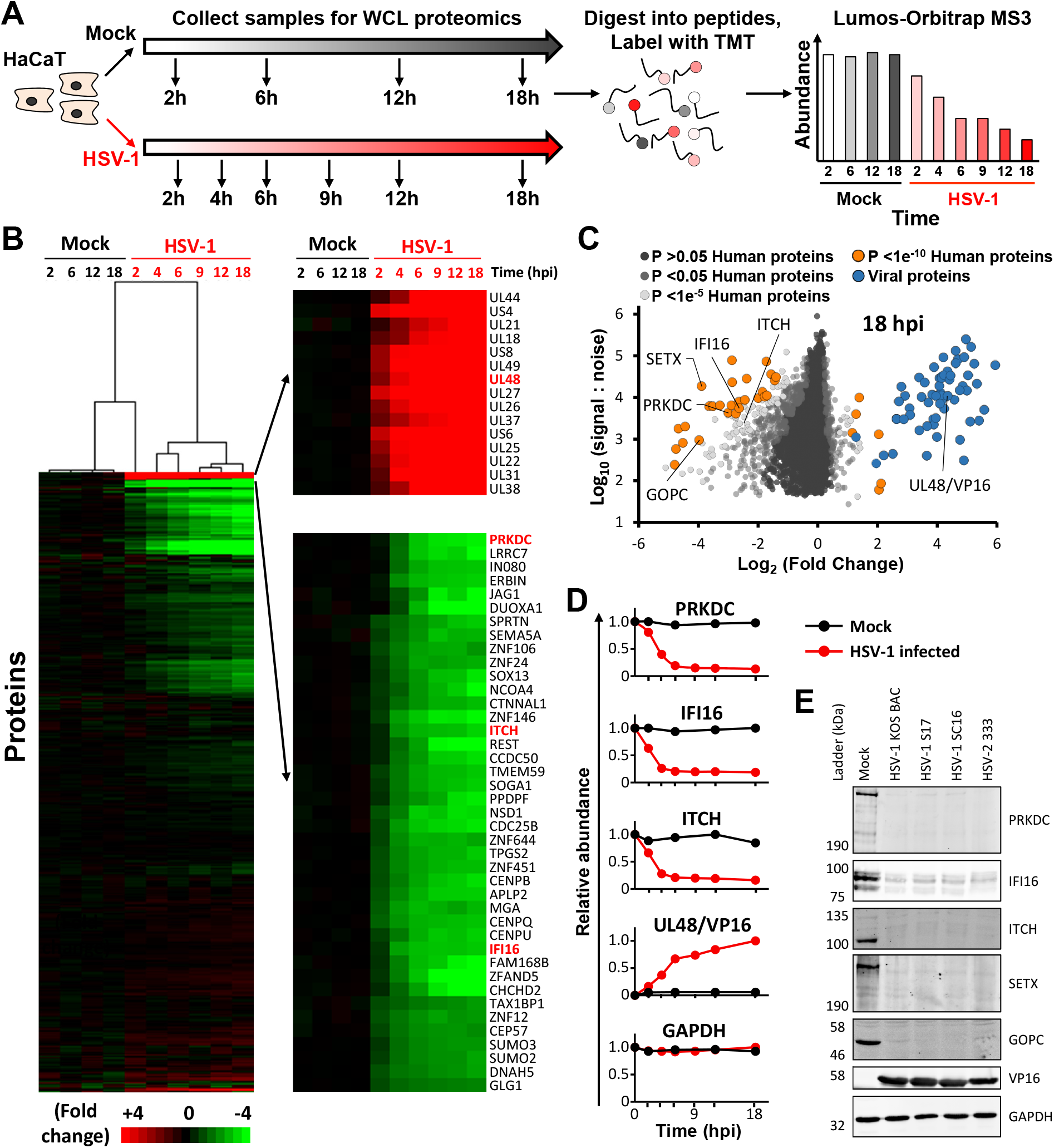
Quantitative temporal analysis of HSV infection. **A**. Schematic of the experimental workflow. HaCaT cells were infected at MOI of 10 or mock infected. Samples were harvested at the stated times and processed for quantitative proteomic analysis. **B**. Hierarchical cluster analysis of all proteins quantified. An enlargement of two subclusters is shown in the right panel, including multiple proteins that were substantially up-or downregulated. **C**. Scatter plot of all proteins quantified at 18 hpi. Fold change is shown in comparison to the average of the mock samples. Benjamini-Hochberg-corrected significance B was used to estimate p-values (Cox and Mann, 2008). **D**. Example temporal profiles for known controls. **E**. Validation of temporal profiles shown in (**D**) by immunoblot of lysates from HaCaT cells infected with a range of HSV strains (MOI of 5 with HSV-1 strains KOS, S17 and SC16, and HSV-2 strain 333).

Temporal analysis of viral protein expression over the whole course of infection can provide a complementary system of protein classification, in addition to enabling direct correlation between viral and cellular protein profiles to give insights into viral-host protein interaction (Soday et al., 2019; Weekes et al., 2014). The number of classes of viral protein expression was determined by clustering viral proteins using the k-means method. This identified at least five distinct temporal protein profiles of viral protein expression (Figure S1, Table S1). Furthermore, by searching data against a 6-frame translation of the HSV-1 strain used (KOS), eight putative new HSV-1 proteins (6FT-ORFs) that increased in abundance over the course of infection were identified (Figure S2A; Table S1).

HSV-1 infection led to >2-fold downregulation of 496 human proteins and >2-fold upregulation of 34 proteins. Mock and immediate early (2h) infection samples clustered separately from early (4, 6h) and late (9, 12, 18h) infection time points. Changes of the greatest magnitude primarily occurred late during infection, as might be expected for a virus with a potent host shutoff activity (Figure 1B). This effect can be observed by a general shift to the left in a scatterplot of fold change (Figure 1C). Multiple host targets known to be specifically downregulated during HSV-1 infection were confirmed, including DNA PKcs (PRKDC) (Lees-Miller et al., 1996; Parkinson et al., 1999), Interferon Gamma Inducible Protein 16 (IFI16) (Orzalli et al., 2012), Promyelocytic Leukemia (PML) (Chelbi-Alix and de The, 1999), Tripartite Motif Containing 27 (TRIM27) (Conwell et al., 2015), Nucleus Accumbens Associated 1 (NACC1) (Sloan et al., 2015), and MORC Family CW-Type Zinc Finger 3 (MORC3) (Sloan et al., 2015) (Figure 1D, Figure 2D and Table S1). Proteomic data was validated by comparison to immunoblot analysis of cells infected for 16 h with three independent strains of HSV-1 and with HSV-2, which suggested that many of the changes observed were conserved phenotypes (Figure 1E). All data are shown in Table S1, in which the ‘‘Plotter” worksheet facilitates interactive generation of temporal graphs of expression of each of the human or viral proteins quantified.

**Figure 2:**
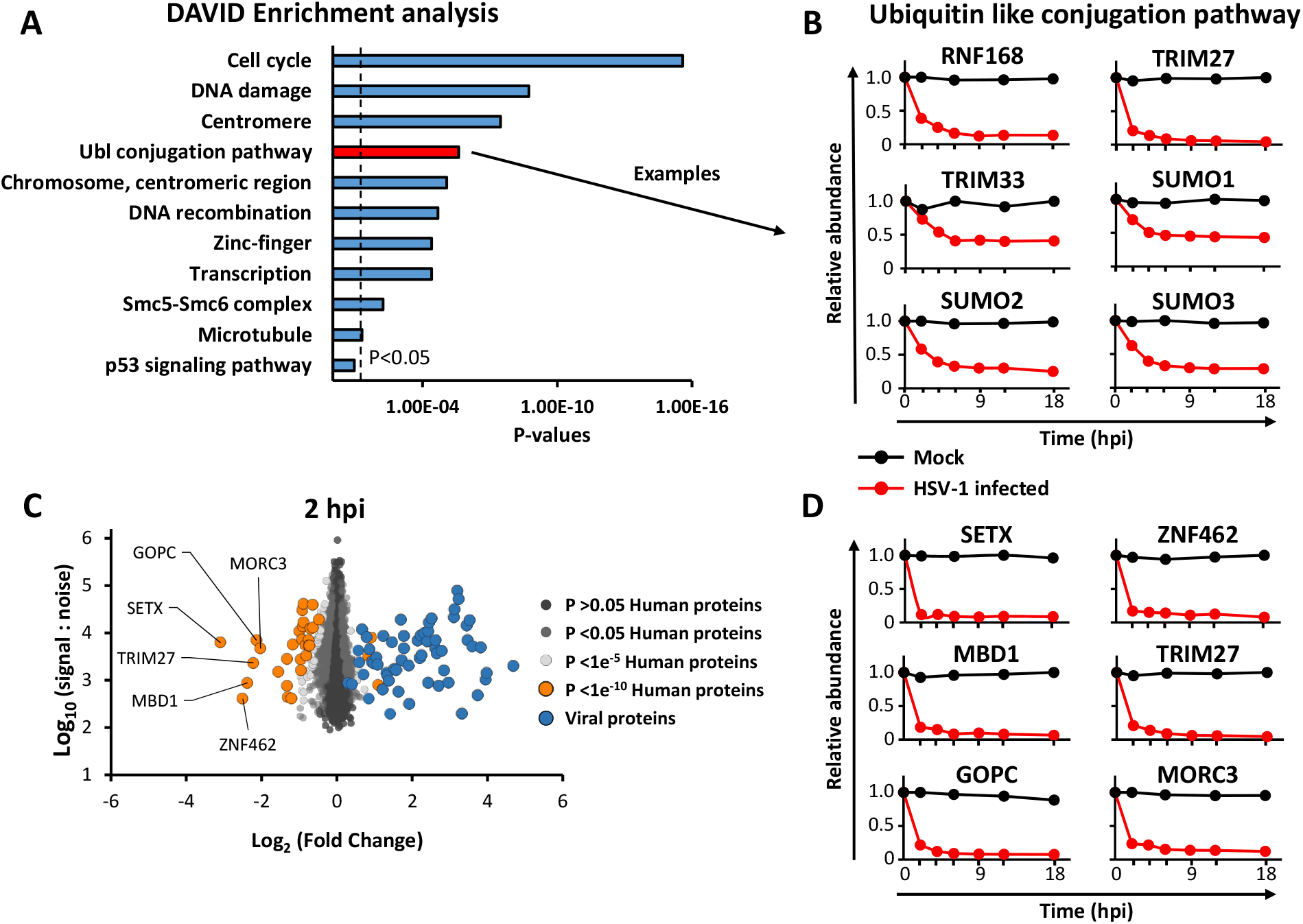
Manipulation of cell host pathways during HSV infection. **A**. DAVID enrichment analysis of all human proteins downregulated >2-fold at any point during infection compared to an average of the four mock samples. A background of all 6956 quantified human proteins was used. Shown are representative terms from each cluster with Benjamini-Hochberg corrected p-values of <0.05. Components of each enriched cluster are shown in Table S2. A similar analysis was performed for proteins upregulated >2 fold, however this did not reveal any significant enrichment. **B**. Example temporal profiles of proteins downregulated from the ubiquitin like (UbL) conjugation pathway. **C**. Scatter plot of all proteins quantified at 2 hpi. Fold change is shown in comparison to the average of the mock samples. Benjamini-Hochberg-corrected significance B was used to estimate p-values (Cox and Mann, 2008). **D**. Temporal profiles of all proteins downregulated during HSV infection >4-fold at 2 hpi.

### Bioinformatic enrichment analysis of HSV-1 infection

DAVID software (Huang da et al., 2009) was used to identify pathways significantly enriched among proteins downregulated >2-fold (Figure 2A). Several of these pathways are known to influence HSV-1 infection, for example cell cycle associated proteins such as cyclin dependent kinases (Schang et al., 1998) and a range of DNA damage response pathways [reviewed in (Smith and Weller, 2015)]. The ubiquitin-like (Ubl) conjugation pathway was significantly enriched, consistent with the known targeting of certain pathway components by herpesviruses to direct cellular prey for degradation. For example, three SUMO family members were downregulated during infection (the fourth was not quantified) (Figure 2B). Components of each enriched cluster are shown in Table S2. A similar analysis of host proteins upregulated >2-fold did not reveal any enriched clusters.

### Identification of host targets most rapidly depleted following HSV-1 infection

Based on the premise that host proteins downregulated early during viral infection are likely to be enriched in factors with antiviral activity, we analyzed proteins downregulated >4-fold at the earliest timepoint after HSV-1 infection (2 hours post infection (hpi); Figure 1C-D). Of the six proteins thus identified, four have previously been shown to be reduced significantly in HSV-1 infected cells (Methyl-CpG Binding Domain Protein 1 (MBD1), MORC3, TRIM27 and Zinc Finger Protein 462 (ZNF462)), of which three were shown to be modulated in an ICP0-dependent manner (MBD1, MORC3, and TRIM27) (Conwell et al., 2015; Sloan et al., 2015). The other two proteins (Senataxin (SETX) and GOPC) have not been previously identified as targets of HSV-1 mediated degradation.

### pUL56 binds NEDD4 family of ubiquitin ligases and GOPC

ITCH, a member of the NEDD4 family of ubiquitin ligases, was rapidly depleted during HSV-1 infection (Figure 1B-D). pUL56 proteins from HSV-1 and HSV-2 interact with ITCH and NEDD4, leading to proteasomal degradation of these targets (Ushijima et al., 2008; Ushijima et al., 2010). pUL56 is a tail-anchored type-II membrane protein found in purified virions (Koshizuka et al., 2002) and contains three PPXY motifs that interact with NEDD4, likely by binding to WW domains (Ushijima et al., 2008). Notably, pUL56 does not contain any lysine residues and is thus likely to be refractory to ubiquitination. To further characterize the cellular binding partners of pUL56, stable isotope labelling of amino acids in cell culture (SILAC) immunoprecipitation–mass spectrometry (IP-MS) analysis was performed using cells expressing GFP-tagged pUL56 or GFP alone (Figures 3A and S3, Table S3). Several members of the NEDD4 family of ubiquitin ligases were enriched in the pUL56 IP, as were multiple Trafficking Protein Particle Complex II (TRAPPCII) subunits. Strikingly, GOPC was also identified as a binding partner of pUL56. Co-precipitation assays demonstrated that the purified GST-tagged pUL56 cytoplasmic domain (residues 1-207) is capable of binding purified GOPC, confirming that these two proteins interact directly (Figure 3B). The N-terminal coiled-coil domain of GOPC mediates its recruitment to the Golgi via an interaction with Golgin-160 (Hicks and Machamer, 2005), whereas the PDZ domain mediates interactions with C-terminal PDZ-binding motifs of cellular partner proteins (Yao et al., 2001). Truncation of GOPC showed that residues 27-236, comprising the N-terminal coiled-coil region, are sufficient to bind to pUL56 (Figure 3C). Immunoprecipitation experiments conducted with cells expressing truncated forms of pUL56 demonstrated that residues 1-157 of pUL56 can mediate efficient binding to GOPC whereas residues 1-104 do not, suggesting that a binding site for GOPC may reside within the 53 amino acid sequence between pUL56 residues 105-157 (Figure 3C). Taken together, these results suggest a model whereby pUL56 binds both GOPC and NEDD4-family of ubiquitin ligases, bringing them in close proximity and thus stimulating the ubiquitination and proteolytic degradation of GOPC.

**Figure 3:**
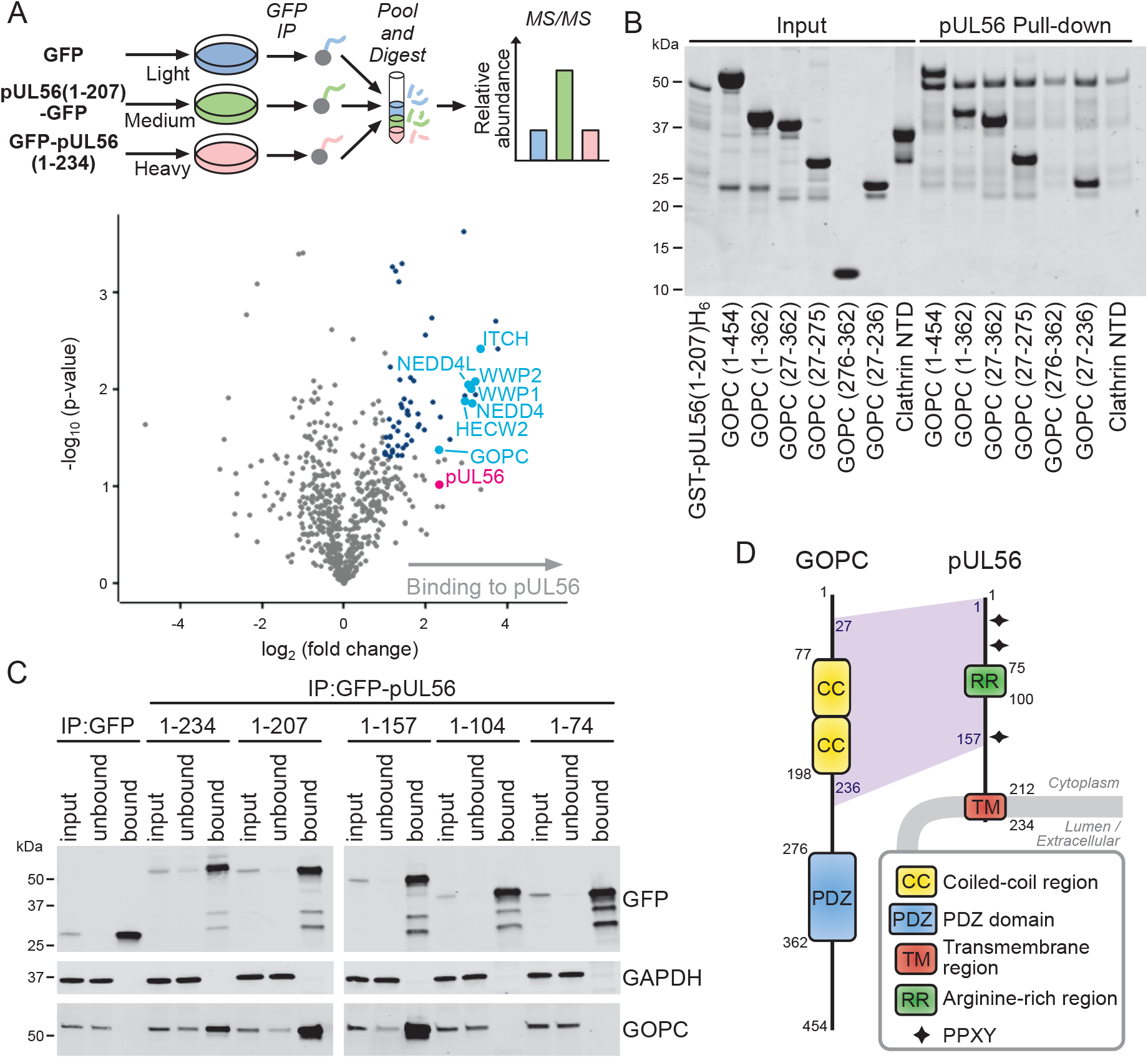
pUL56 binds GOPC and cellular ubiquitin ligases. **A**. SILAC-labelled HEK293T cells were transfected with GFP-tagged pUL56 cytoplasmic domain (residues 1-207) or GFP alone and subjected to immunoprecipitation (IP) using a GFP affinity resin. In the volcano plot, the horizontal-axis shows average fold enrichment in IP of pUL56(1-207)-GFP compared to GFP across three biological replicates and the vertical axis shows significance (two sided t-test) across the three replicates. Significantly enriched proteins (>2-fold enrichment and p < 0.05) are colored blue and selected proteins are annotated. **B**. Pull-down experiment using purified recombinant components, demonstrating that the GST-tagged pUL56 cytoplasmic domain interacts directly with the coiled-coil region of GOPC. The peptide-binding N-terminal domain of clathrin heavy chain (Clathrin NTD) and GST were used as control prey and bait proteins, respectively. **C**. Co-IP of GOPC with GFP-tagged pUL56 and truncations thereof. Immunoblots were stained with the antibodies shown. **D**. Schematic representation of pUL56 and GOPC.

### pUL56 mediates degradation of GOPC via the proteasome

To identify the mechanism of GOPC degradation, cells were infected with wild-type (WT) HSV-1 or HSV-1 lacking expression of pUL56 (ΔUL56). Viruses lacking expression of the viral proteins ICP0 (ΔICP0) or vhs (Δvhs) were also included, as both are known to deplete host proteins. Cells were further treated with or without the proteasomal inhibitor MG132. GOPC was degraded during HSV-1 infection in a pUL56-dependent and MG132-inhibitable fashion, whereas GOPC degradation was independent of both ICP0 and vhs (Figure 4A-B). Expression of pUL56 by transfection or in an inducible cell line demonstrated that this protein is sufficient for GOPC degradation in the absence of other HSV-1 factors (Figures 4C-D). HSV-1 pUL56 contains three PPXY motifs, which mediate interaction with NEDD4 family of E3 ubiquitin ligases (Ushijima et al., 2010). To test the importance of these PPXY motifs in pUL56 for GOPC degradation a recombinant virus was generated in which all three motifs were mutated to AAXA. This triple mutant phenocopied the deletion virus, failing to degrade GOPC even though pUL56 expression was maintained (Figure 4E). Overall, these data suggest that pUL56 recruits NEDD4 family ubiquitin ligases to mediate the proteasomal degradation of GOPC.

**Figure 4:**
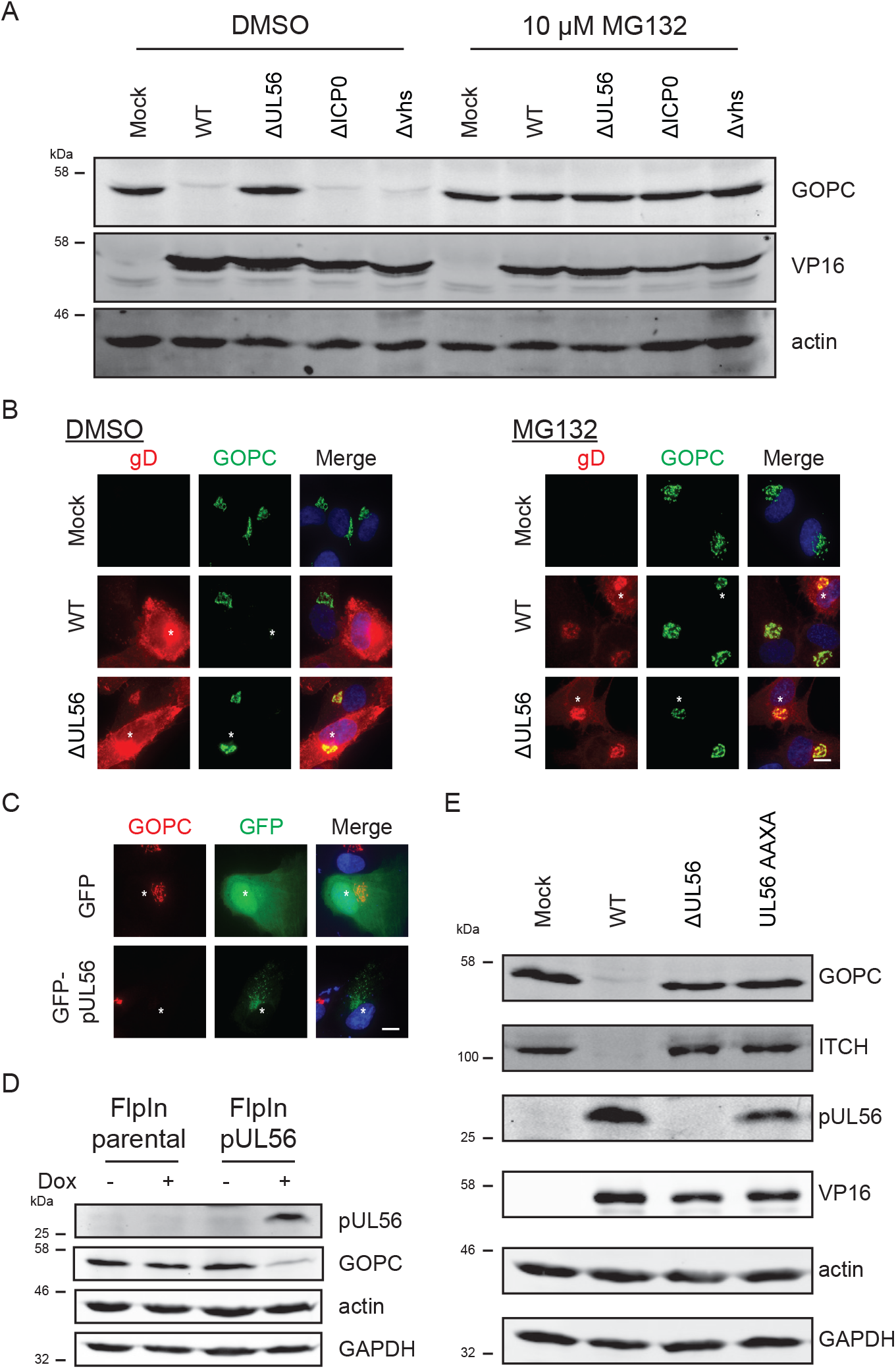
pUL56 is necessary and sufficient for GOPC degradation. **A**. HaCaT cells were infected at MOI of 10 with the indicated virus. After 2 h, media was replaced with 10 μM MG132 or carrier (DMSO) in DMEM for the remainder of the infection. Cell lysates were harvested 16 hpi. **B.** HFF hTERT cells were infected at MOI of 1 then treated with MG132 or carrier as described in (**A**). At 6 hpi, samples were fixed and stained for GOPC (green) and the infection control gD (red). The merge includes DAPI (blue) and the scale bar represents 10 μm. **C.** HFF cells were transfected with pUL56-GFP. One day post-transfection cells were fixed and stained for GOPC (red). The merge includes DAPI (blue) and the scale bar represents 10 μm. **D.** Expression of pUL56 was induced with doxycycline in a clonal 293 FlpIn cell line. Cell lysates were harvested 1 day after induction. **E.** HaCaT cells were infected at MOI of 10 with the indicated virus and cell lysates were harvested 16 hpi.

### Replication of HSV-1 in cell culture is independent of pUL56

The rapid depletion of GOPC from cells during HSV-1 infection implies that removal of this host protein may be important for efficient viral replication. However, HSV-1 ΔUL56, where endogenous levels of GOPC are maintained, demonstrated effectively identical growth kinetics to HSV-1 WT (Figure 5A). Plaque size analysis also demonstrated no defects in cell-to-cell spread for HSV-1 ΔUL56 compared to WT (Figure 5B-C). These data demonstrate that pUL56 is dispensable for HSV replication in cell culture, consistent with previous reports (Ushijima et al., 2008). Given that viruses do not usually retain genes of no benefit, this suggests that pUL56 plays a role during viral replication *in vivo*, perhaps during establishment, maintenance, or reactivation from latency. Alternatively, pUL56 may be a virulence factor involved in modulating antiviral immune responses against HSV-1, as is the case for a number of herpesvirus proteins that are dispensable in cell culture but important for replication *in vivo*, for example vhs (Strelow and Leib, 1995).

**Figure 5:**
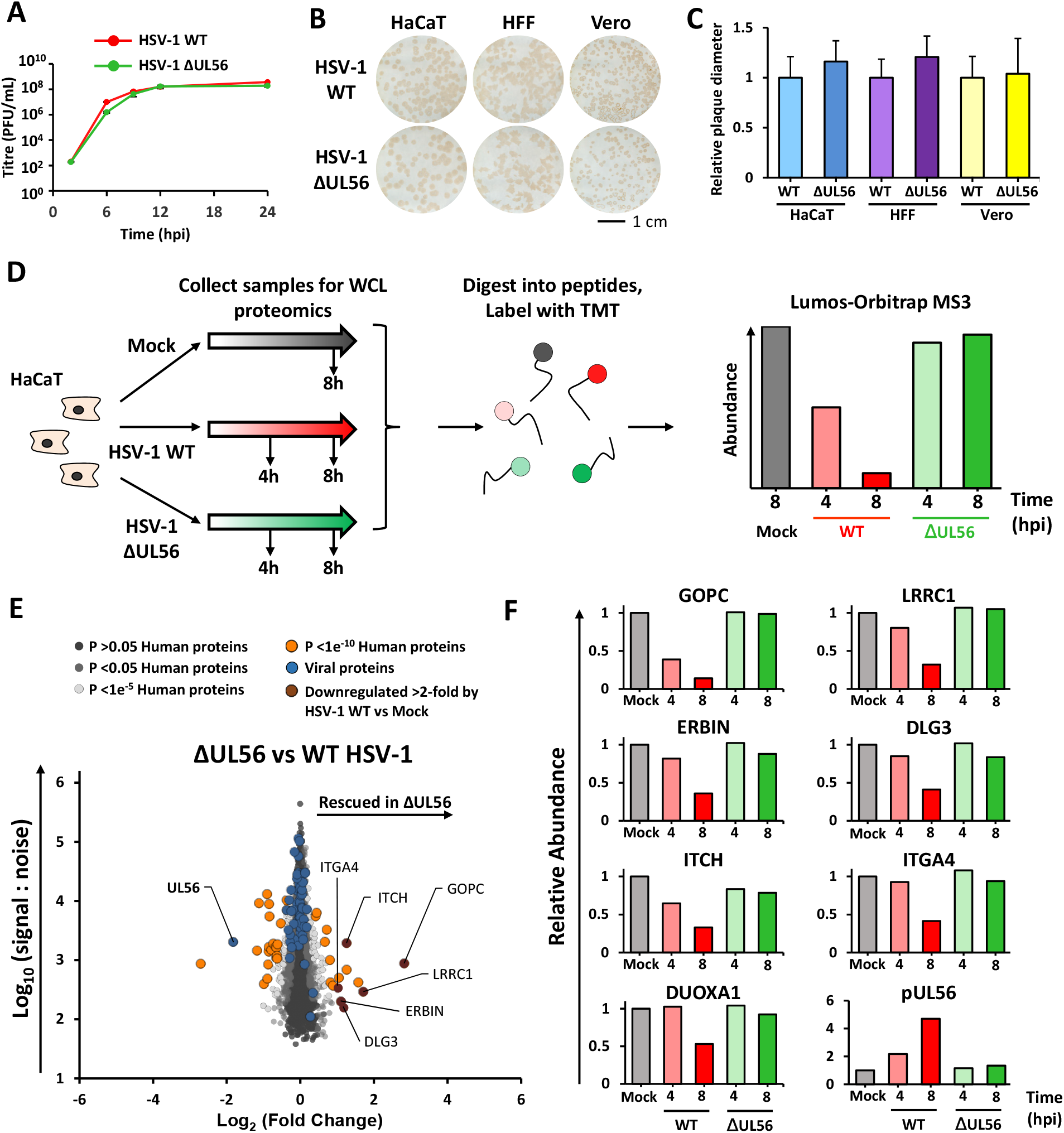
Identification of pUL56 degradation targets. HaCaT cells were infected at MOI of 10 in biological duplicate. Error bars: +/-standard deviation (SD). Plaque assays of HSV-1 WT and HSV-1 ΔUL56 in HaCaT, HFF hTERT, and Vero cells in biological duplicate. Cells were subsequently immunostained for the viral glycoprotein gD. **C**. Plaque diameters from (**B**) were measured and normalized to HSV-1 WT. Error bars: +/- SD, n = 35 to 67. **D**. Schematic of the proteomics workflow. Cells were infected at MOI of 10 or mock infected. **E**. Scatter plot of all proteins quantified, comparing HSV-1 WT and HSV-1 ΔUL56 at 8 hpi. Benjamini-Hochberg-corrected significance B was used to estimate p-values (Cox and Mann, 2008). **F**. Temporal profiles of all proteins downregulated >2-fold by HSV-1 WT vs mock and additionally rescued >2-fold by HSV-1 ΔUL56.

### Identification of host proteins specifically depleted by pUL56

ICP0 and vhs are known to cause extensive remodelling of host protein expression to facilitate viral replication (Boutell and Everett, 2013; Smiley, 2004). Our data now suggest that pUL56 also contributes to host protein depletion but in a more targeted manner. To identify cellular proteins depleted by pUL56, HaCaT cells were infected with HSV-1 WT or ΔUL56 and analyzed by TMT-based proteomics (Figure 5D, Table S4). Of the 7696 human proteins quantified, only a small number exhibited significant abundance changes between the WT and ΔUL56 infections, and the largest change observed was for GOPC (Figure 5E-F). A small number of other potential targets of pUL56 were identified, defined by >2-fold reduced abundance in HSV-1 WT samples compared to mock and ΔUL56 samples. These included Discs Large MAGUK Scaffold Protein 3 (DLG3), Leucine Rich Repeat Containing 1 (LRRC1), and Erbb2 Interacting Protein (ERBIN), which may function as a complex: both LRRC1 (aka LANO) and ERBIN have been shown to interact with DLG proteins (Saito et al., 2001). The discs-large (DLG) family have a number of proposed functions including regulation of cell polarity and tight junction formation, and they are targeted for degradation by a number of viral families (Kong et al., 2014; Lee et al., 1997; Roberts et al., 2012). Remodelling cell polarity through pUL56-mediated degradation of these host proteins may facilitate HSV-1 spread *in vivo*.

Searching this TMT dataset against a 6-frame translation of KOS-strain HSV-1 identified 14 novel HSV-1 proteins that increased in abundance over the course of infection, including 9 that were not identified in the initial QTV experiment (Figure S2B, Table S4).

### pUL56-activity alters the plasma membrane proteome

Modulation of proteins at the cell surface is an immune evasion strategy utilized by multiple viruses. Since GOPC regulates the trafficking of certain proteins to the plasma membrane (Cheng et al., 2002), destruction of GOPC through the activity of pUL56 may be a mechanism to specifically modify the surface presentation of proteins in HSV-1 infected cells. Plasma membrane profiling was thus performed on cells infected with HSV-1 WT or ΔUL56 at 6 hpi using SILAC-based mass spectrometry (Figure 6, Table S5). Hierarchal clustering of the resulting data identified host proteins that are less abundant at the plasma membrane of HSV-1 WT infected cells and rescued by pUL56 deletion. These included immune signalling proteins TLR2 and IL18R1 as well as DUOX1 and several members of the solute carrier (SLC) family of proteins. TLR2 is a pattern recognition receptor that has a well-established activity against bacterial pathogen-associated molecular patterns (PAMPs), but also recognizes HSV-1 and HCMV glycoproteins (Boehme et al., 2006; Cai et al., 2013; Leoni et al., 2012). In response to herpesvirus infection, TLR2 plays a role in inducing interferon γ in neurons, cytokines in peritoneal macrophages, as well as controlling viral load in the CNS (Lima et al., 2010; Sorensen et al., 2008). IL18 is a proinflammatory cytokine that binds IL18R1, which is important for innate immune responses to HSV-2 infection *in vivo* (Harandi et al., 2001). Downregulating these immune receptors from the cell surface may be a pro-viral strategy to decrease inflammation and immune activation. DUOX1 (dual oxidase 1) is a transmembrane protein that can generate H_2_O_2_ and functions in lactoperoxidase-mediated antimicrobial defence at mucosal surfaces (Sarr et al., 2018). Production of H_2_O_2_ has been shown to inhibit the splicing of influenza A virus (IAV) transcripts and decrease production of infectious virus, and IAV has been shown to downregulate DUOX1 (Strengert et al., 2014). Removing DUOX1 from the plasma membrane may be similarly proviral for HSV-1 by inhibiting H_2_O_2_ production. The mechanism by which HSV-depletes DUOX1 from the plasma membrane may be through pUL56-dependent degradation of DUOXA1 (Figure 5F) as DUOXA1 is a chaperone required for the maturation and transport of DUOX1 from the ER to the plasma membrane (Grasberger and Refetoff, 2006).

**Figure 6:**
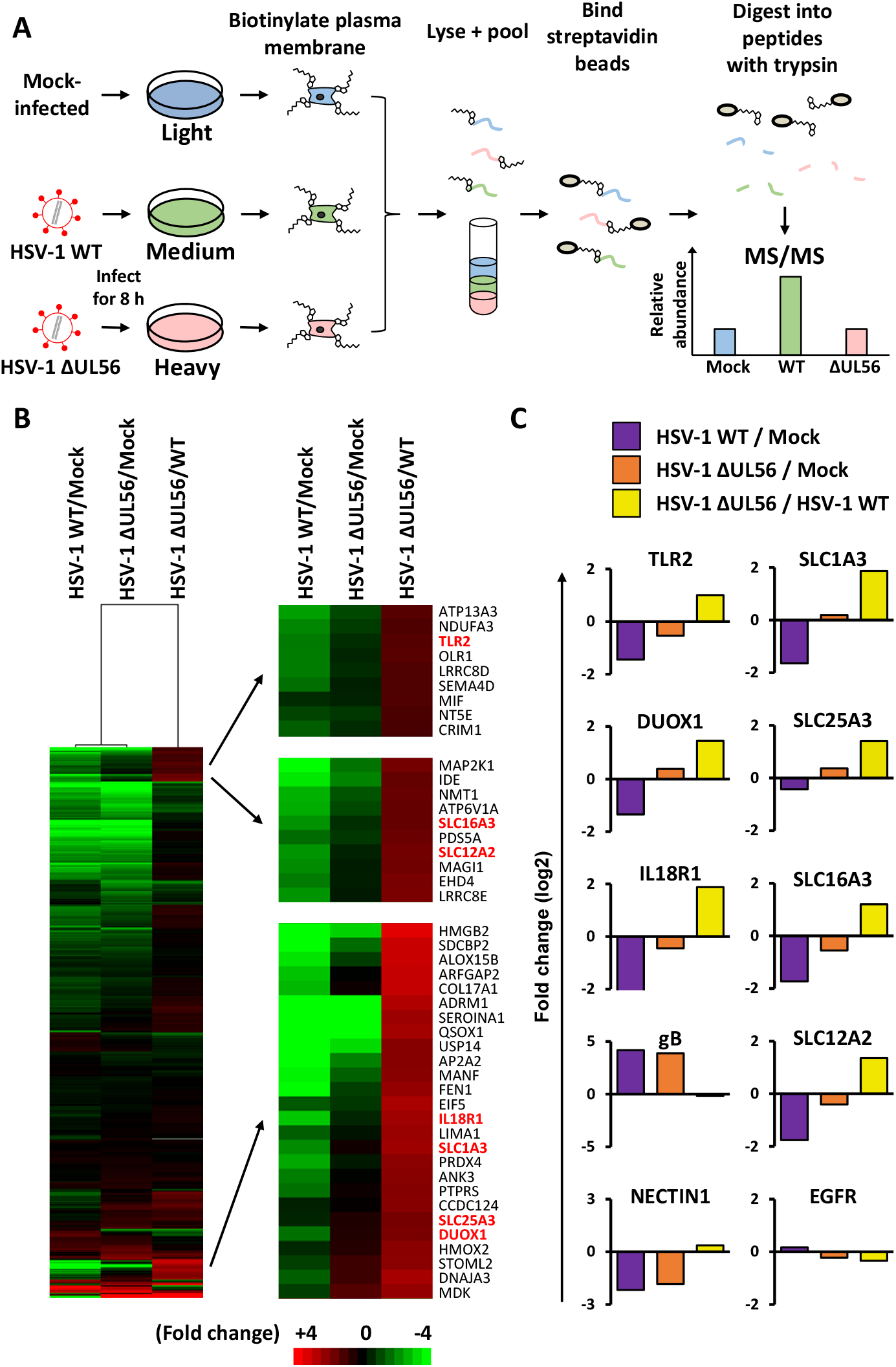
pUL56 modulates immune receptors through control of host trafficking to the plasma membrane. **A**. Schematic of the experimental workflow. SILAC labelled cells were infected at MOI of 10 or mock infected. Samples were harvested 6 hpi and processed for plasma membrane enrichment and subsequent quantitative mass spectrometry. **B**. Hierarchical cluster analysis of fold change values for each pair-wise comparison. Proteins were included if they are annotated as plasma membrane (PM), cell surface (CS), or extracellular (XC) by Gene Ontology (GO), or with a short GO term as previously described (Weekes et al., 2014). An enlargement of three subclusters is shown in the right panel, which included proteins downregulated during infection with HSV-1 WT but rescued by infection with HSV-1 ΔUL56. **C**. Profiles of example proteins that were downregulated >2-fold by HSV-1 WT and rescued >2-fold by HSV-1 ΔUL56 are shown as well as controls.

### TLR2 signalling is prevented by pUL56

To determine if loss of TLR2 from the cell surface was due to disruption of GOPC mediated trafficking, we generated GOPC knockout HaCaT cells using CRISPR/Cas9 gene engineering. Three independent single-cell clones generated from two independent sgRNAs targeting GOPC were stained for surface TLR2 and analyzed by flow cytometry. Normal HaCaT cells included a TLR2^+^ population, whereas all three GOPC knockout clones exhibited reduced cell surface TLR2 (Figure 7A-B), suggesting TLR2 surface expression is GOPC-dependent.

**Figure 7:**
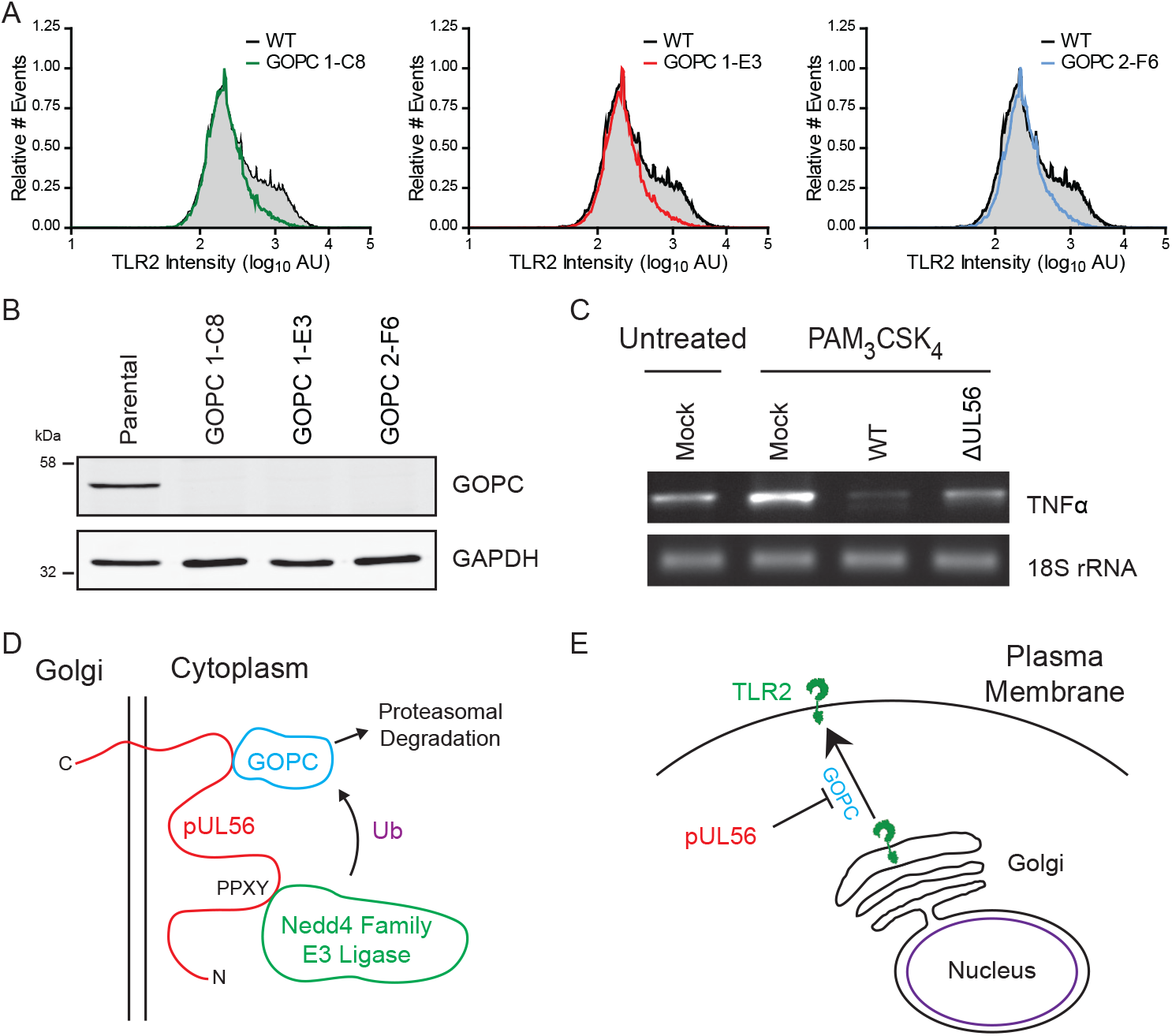
pUL56 degrades GOPC to prevent presentation of TLR2 at the plasma membrane. **A**. Flow cytometry analysis of TLR2 levels at the plasma membrane of HaCaT wild-type (WT) cells, and three GOPC knockout (GOPC) clones. Single cell clones (C8, E3, and F6) were isolated from CRISPR knockout cells made from 2 independent gRNAs (GOPC 1 and GOPC 2). **B**. Immunoblot analysis of GOPC knockout cells. **C**. Differentiated THP1 cells were infected at MOI of 10 with the indicated virus and at 6 hpi 100 ng/mL PAM3CSK4 was added. At 1 day post infection, RNA was extracted and analyzed for TNFα mRNA relative to 18S rRNA. **D**. Model illustrating how pUL56 mediates the degradation of GOPC through recruitment of host Nedd4 family E3 ubiquitin ligases via PPXY motifs. **E**. Degradation of GOPC results in the loss of specific anti-viral proteins, such as TLR2, from the cell surface.

Human monocytes respond to HSV-1 infection via TLR2 (Ahmad et al., 2008). To assess the biological effect of TLR2 downregulation during infection, receptor activity was measured in the THP1 human monocytic cell line. Cells were infected with HSV-1 WT or ΔUL56 and then stimulated with the TLR2 agonist PAM_3_CSK_4_. The expression levels of TNFα mRNA, which is stimulated by TLR2 signalling, were analyzed by RT-PCR. PAM_3_CSK_4_ treatment of mock infected cells stimulated TNFα expression as expected, and infection with HSV-1 WT supressed TNFα levels to below unstimulated controls. Cells infected with HSV-1 ΔUL56 showed greater levels of TNFα expression relative to HSV-1 WT, although the levels were still suppressed compared to mock infected cells stimulated with PAM_3_CSK_4_ (Figure 7C). This suggests that the functional consequences of pUL56-stimulated degradation of GOPC includes inhibition of TLR2 signalling.

## Discussion

In this study we combined three powerful unbiased proteomic analysis techniques, quantitative temporal viromics (Figures 1, 2 and 5), affinity enrichment (Figure 3) and plasma-membrane proteomics (Figure 6), to identify that HSV-1 protein pUL56 promotes degradation of the host-cell trafficking factor GOPC and in doing so lowers the abundance of important immune signalling molecules at the plasma membrane of infected cells. Biochemistry and cell biology experiments (Figures 3, 4, 5 and 7) confirmed that pUL56 binds directly to GOPC, is both necessary and sufficient to promote GOPC degradation, requires the recruitment of the NEDD4 family of ubiquitin ligases via its PPXY motifs for such degradation, and results in changes in the cell-surface proteome through the loss of GOPC. The proteomic datasets presented in this manuscript represent a rich resource for identifying and characterizing the mechanisms by which herpesviruses modulate both the whole-cell and plasma-membrane proteomes of infected cells.

### Temporal insights into HSV-1 infection

The QTV data presented herein represent the most comprehensive analysis of host-cell proteome changes upon HSV-1 infection to date, with almost 7000 host proteins quantified, and provide important insights into the kinetics of HSV-1 viral protein production. K-means analysis identified five distinct profiles of protein expression (Figure S1). Immediate early and early genes were found in the same class (Tp2). This presumably arises from the high multiplicity of infection, required for synchronous and complete infection, and the use of 2 hpi as the earliest time point. These conditions may have masked some of the differences in the kinetic profiles of immediate early and early gene classes. Interestingly, late genes appeared to cluster into three distinct groups (Tp3-5). While late genes have previously been divided into late and true late classes, dependent on the requirement for prior genome replication (Kibler et al., 1991), our data suggests that an intermediate kinetic class may exist. Alternatively, this data may highlight that viral proteins can mature at different rates despite their expression being induced at the same time.

This kinetic analysis of HSV-1 protein abundance identified that ICP47 (US12) has a unique temporal profile (Tp1, Figure S1). Unlike all other viral proteins, where the abundance increases throughout infection, the amount of ICP47 peaks early during infection and the protein is subsequently downregulated. ICP47 binds and inhibits the MHC class I peptide loading complex Transporter of Antigenic Peptides (TAP), preventing peptide presentation at the cell surface and promoting immune evasion (Hill et al., 1995). The varying abundance of ICP47 during infection might therefore have the effect of balancing evasion of CD8+ T-cells whilst preventing activation of NK cell killing, by precisely regulating the level of MHC-I reduction at the cell surface.

### HSV-1 pUL56 degrades GOPC by recruiting cellular E3 ligases

HSV-1 strains lacking pUL56 are attenuated in animal models (Berkowitz et al., 1994; Kulej et al., 2017; Rosen-Wolff et al., 1991), despite the protein being dispensable for virus replication in cultured cells (Figure 5A-C) (Ushijima et al., 2008). Our data provide a molecular mechanism by which pUL56 may enhance virulence during infection, by promoting the degradation of GOPC and subsequent down-regulation of immune signalling molecules from the surface of infected cells.

Previous studies from HSV-1 and HSV-2 have shown pUL56 interact with ITCH and NEDD4, leading to their degradation, but the importance of this activity remained elusive (Ushijima et al., 2008; Ushijima et al., 2010). Our IP-MS data revealed that pUL56 binds multiple cellular NEDD4-family ubiquitin ligases and the trafficking factor GOPC (Figure 3A). We show that pUL56 binds directly to the coiled-coil region of GOPC (Figure 3B), is necessary for the proteasome-mediated degradation of GOPC in HSV-1 infected cells (Figure 4A-B), and is sufficient to promote GOPC degradation in the absence of infection (Figure 4C-D). Furthermore, we show that the NEDD4-binding PPXY motifs of pUL56 are required for GOPC degradation (Figure 4E). Taken together, these data strongly support a model whereby pUL56 binds simultaneously to GOPC and the NEDD4 family of ubiquitin ligases in order to promote GOPC ubiquitination and proteasomal degradation (Figure 7D). pUL56 is itself protected from degradation as it does not contain lysine residues to which ubiquitin could be conjugated.

GOPC is rapidly degraded during HSV-1 WT infection (Figure 2D) and its abundance is restored during infection with HSV-1 ΔUL56 (Figure 5E). Several other proteins are also rescued when comparing HSV-1 WT to ΔUL56 infection. This may reflect direct pUL56-mediated degradation or be an indirect consequence caused by the loss of GOPC.

### HSV-1 degrades a trafficking factor to modify the surface of infected cells

Many viruses modify the surface of infected cells in order to modulate the host immune response. For example, HIV-1 Vpu recruits an E3 ligase to promote the ubiquitination and the degradation of several cell-surface proteins (Matheson et al., 2015). Alternatively, it has been shown that multiple proteins from human cytomegalovirus act via distinct mechanisms to restrict the cell-surface presentation of class I MHC and NK cell receptors (Wilkinson et al., 2008). Using global unbiased approaches, we have now identified that HSV-1 pUL56 modifies the surface abundance of cellular immune signalling proteins by specifically degrading a key cellular trafficking factor. This factor, GOPC, may be a common target for modulation by viruses: human papillomavirus type 16 E6 protein was shown to bind GOPC and mediate its degradation through the host E3 ubiquitin ligase E6AP (Jeong et al., 2007). Unlike pUL56, E6 binds to the PDZ domain of GOPC through a PDZ-binding motif. In addition, the classical swine fever virus NS2 protein bound GOPC in a yeast two-hybrid screen (Kang et al., 2012), although it has not yet been determined if GOPC is degraded during infection with this virus.

The pUL56 homologues from equine herpesvirus type 1 (EHV-1) and type 4 (EHV-4) share only 20% identity with HSV-1 pUL56, yet both are type II transmembrane proteins that possess multiple PPXY motifs and have few or no cytoplasmic lysine residues. Interestingly, both EHV-1 and EHV-4 have been shown to downregulate MHC-1 from the surface of infected cells in a pUL56-dependent fashion (Ma et al., 2012; Said et al., 2012). Similarly, U24 from human herpesvirus 6A (HHV-6A) is a tail-anchored (type-II) membrane protein containing a PPXY motif and has been shown to downregulate the T-cell receptor (Koshizuka et al., 2018; Sullivan and Coscoy, 2008). It therefore seems likely that EHV pUL56 and HHV-6A U24 also recruit NEDD4-family ubiquitin ligases to degrade specific cellular proteins and thus modify cell-surface protein abundance, but the direct targets of these proteins remain unknown.

In conclusion, our data have identified that HSV-1 pUL56 targets GOPC for proteasomal degradation, thereby removing immune signalling molecules from the plasma membrane. This represents an elegant and efficient mechanism by which HSV-1 can remodel the surface of infected cells. The degradation of GOPC by other viruses such as human papillomavirus suggests that targeting of GOPC specifically, or trafficking factors more generally, may represent a common mechanism by which viruses modulate the host-cell surface to evade host immune surveillance.

## Acknowledgments

We thank Kate Heesom (University of Bristol) for SILAC immunoprecipitation mass spectrometry sample processing and data acquisition. We thank Steve West (Frances Crick Institute) for the SETX antibody, Nick Gay and Atul Udgata (University of Cambridge) for THP1 cells, Susanna Colaco for superb technical assistance and Janet Deane for helpful discussions. This work was funded by a Wellcome Trust PhD studentships to J.M., C.S. and H.G.B., a Sir Henry Dale Fellowship, jointly funded by the Wellcome Trust and the Royal Society (098406/Z/12/B), to S.C.G., a Wellcome Trust Senior Clinical Research Fellowship (108070/Z/15/Z) to M.P.W. and a BBSRC Research Grant (BB/M021424/1) to C.M.C. This research was supported by the Cambridge NIHR BRC Cell Phenotyping Hub.

## Author Contributions

Conceptualization, T.K.S., J.M., S.C.G., M.P.W., C.M.C.

Methodology, T.K.S., C.T.R.D., E.E., S.C.G., M.P.W., C.M.C.

Validation, T.K.S., C.T.R.D., J.M., C.R.B.

Investigation, T.K.S., C.T.R.D., J.M., V.C., C.R.B., H.G.B., C.S., E.E.

Data Curation T.K.S., C.T.R.D., J.M., S.C.G., M.P.W., C.M.C.

Writing – Original Draft, T.K.S., C.T.R.D., J.M.

Writing – Review & Editing T.K.S., C.T.R.D., J.M., S.C.G., M.P.W., C.M.C.

Visualization, T.K.S.; C.T.R.D., J.M., S.C.G., M.P.W., C.M.C.

Supervision S.C.G., M.P.W., C.M.C.

Project Administration C.M.C.

Funding Acquisition S.C.G., M.P.W., C.M.C.

## Supplementary Table Legends

Table S1, related to Figure 1: Quantitative temporal analysis of HSV infection.

Interactive spreadsheet of the quantitative temporal viromics data from the HSV-1 whole cell lysate time course. The “Data” worksheet shows minimally annotated protein data, with only formatting and normalization modifying the raw data. The “Plotter” worksheet enables generation of individual protein abundance changes from the time course. The ‘Viral classes’ tab shows the K-means clusters of all canonical HSV-1 proteins, a list of 5 clusters are shown. The ‘MS Quantification’ tab shows the number of proteins and peptides quantified. The ‘Novel 6FT-ORFs’ tab contains details of putative new HSV-1 proteins that increased in abundance over the course of infection.

Table S2, related to Figure 2: Manipulation of cell host pathways during HSV infection.

DAVID functional enrichment analysis from proteins downregulated >2-fold against a background of all proteins quantified. Only significant (Benjamini-Hochberg corrected) clusters are shown. There were no significant clusters amongst proteins upregulated >2-fold.

Table S3, related to Figure 3: Identification of cellular interaction partners of pUL56.

Spreadsheet listing the SILAC ratios and statistical analysis of proteins quantified in pull-downs of pUL56 followed by mass spectrometry (IP-MS). Two different constructs of pUL56 encompassing either the full-length protein or its cytoplasmic domain were tested and the respective results are listed in separate tabs.

Table S4, related to Figure 5: Identification of pUL56 degradation targets.

Interactive spreadsheet displaying whole cell protein changes between cells infected with HSV-1 WT, HSV-1 ΔUL56 or mock. The “Data” worksheet shows minimally annotated protein data, with only formatting and normalization modifying the raw data. The “Plotter” worksheet enables generation of individual protein abundance changes, comparing the different viruses and time points. The ‘MS Quantification’ tab shows the number of proteins and peptides quantified. The ‘Novel 6FT-ORFs’ tab contains details of putative new HSV-1 proteins that increased in abundance over the course of infection.

Table S5, related to Figure 6: pUL56 modulates immune receptors through control of host trafficking to the plasma membrane.

Interactive spreadsheet of plasma membrane protein changes between cells infected with HSV-1 WT, HSV-1 ΔUL56 or mock. The “Data” worksheet shows minimally annotated protein data, with formatting and normalization modifying the raw data. GO terms were used to identify proteins associated with the plasma membrane, as described in the text. The “Plotter” worksheet enables generation of individual protein ratios between the three conditions. The ‘MS Quantification’ tab shows the number of proteins and peptides quantified.

## Methods

### Contact for Reagent and Resource Sharing

Further information and requests for resources and reagents should be directed to and will be fulfilled by the Lead Contact, Colin M. Crump.

#### Mammalian cell culture

HaCaT (Boukamp et al., 1988), telomerase immortalized human foreskin fibroblasts HFF hTERT (McSharry et al., 2001), Vero (ATCC), HEK293T (ATCC), and Flp-In™-293 (ThermoFisher) cells were maintained in Dulbecco’s Modified Eagle’s Medium (DMEM). THP1 cells were grown in RPMI 1640. All medium was supplemented with 10% fetal bovine serum (FBS), 2 mM L-glutamine, 100 U/mL penicillin, and 100 μg/mL streptomycin and cells were grown at 37°C in a humidified 5% CO_2_ atmosphere. HFF hTERT, HaCaT, and THP1 cells are male; no gender is provided for the other cell lines. For stable isotope labelling of amino acids in cell culture (SILAC) experiments, HEK 293T cells were grown in SILAC medium (high glucose DMEM lacking arginine and lysine, Life Technologies) supplemented with 10% dialyzed fetal bovine serum (10 kDa cutoff), 2 mM glutamine, 100 U/mL penicillin and 100 µg/mL streptomycin. Media were supplemented with 84 mg/L arginine (light, unlabelled; medium, Arg6 (^13^C6); heavy, Arg10 (^13^C6, ^15^N4)) and 146 mg/L lysine (light, unlabelled; medium, Lys4 (^2^H4); heavy, Lys8 (^13^C6, ^15^N2)). Cells were maintained in SILAC media for at least five passages before use to ensure complete labelling.

THP1 cells (10^6^ cells/mL) were differentiated by resuspension in RPMI 1640 supplemented with in 16 nM phorbol 12-myristate-13-acetate (PMA) and seeded at 2.5×10^5^ cells/cm^2^. After treatment with PMA for 1 day, the media was changed to RPMI without PMA and the cells were given a recovery period of 2 days before being infected.

#### Viruses

All HSV-1 strain KOS viruses were reconstituted from a bacterial artificial chromosome (Gierasch et al., 2006). The deletion mutants were generated by inserting three tandem stop codons in frame using the two-step Red recombination method (Tischer et al., 2010). For ΔUL56 this is after residue 21, for ΔICP0 this is after residue 11, and for Δvhs this is after residue 45 (Zenner et al., 2013). HSV-1 S17 and HSV-2 333 were from S. Efstathiou (University of Cambridge), and HSV-1 SC16 was from A. Minson (University of Cambridge). Crude stocks were generated by infecting Vero cells at MOI of 0.01. After 3 days, the cells were scraped and isolated by centrifugation at 900×g for 5 min. They were resuspended in 1 mL of complete media per T150 used and freeze/thawed thrice at −70°C before being aliquoted, titred, and stored at −70°C until required.

#### Gradient purification of HSV-1

HaCaT cells were seeded and infected with crude virus stocks at MOI of 0.1. After 2 days, the cells were scraped and the cell debris was removed by centrifugation at 900×g for 5 min. The supernatant was ultracentrifuged at 24,000×g for 1.5 h, and the pellet was resuspended in 1% FBS in PBS on ice overnight. This solution was overlaid on a 5-15% (w/v) continuous Ficoll in PBS gradient and ultracentrifuged at 17,500×g for 1.5 h. The virion band was isolated via side-puncture. This solution was diluted 10-fold in PBS, and the virus was pelleted by ultracentrifugation at 49,000×g for 2 h. The pellet was resuspended in PBS on ice overnight. This solution was aliquoted, titred, and stored at −70°C until required.

#### Antibodies

Primary antibodies used were: GOPC (Millipore, MABC731), DNA PKcs (Santa Cruz Biotechnology, sc-5282), IFI16 (Santa Cruz Biotechnology, sc-8023), SETX (OY7) (Yuce and West, 2013), ITCH (Santa Cruz Biotechnology, sc-28367), WWP2 (Santa Cruz Biotechnology, sc-398090), GAPDH (ThermoFisher Scientific, AM4300), Actin (Abcam, AC-40), TLR2 (BioLegend, 153003), GFP (Clontech, JL-8), gD (LP2) (Minson et al., 1986), VP16 (LP1; Abcam, ab110226) (McLean et al., 1982), pUL56 (see below). Secondary antibodies used were: Alexa Fluor 488 labelled Donkey anti-Rabbit IgG (ThermoFisher Scientific, A21206), Alexa Fluor 568 labelled Donkey anti-Mouse IgG (ThermoFisher Scientific, A10037), IRDye 680LT Goat anti-Mouse IgG (LiCor, 926-68020), IRDye 800CW Donkey anti-Rabbit IgG (LiCor, 926-32213), Goat anti-Mouse HRP conjugate (CiteAb, P0447).

An antibody against pUL56 was generated by commercial immunization of a rabbit using two peptides (peptide 1: NH_2_-CTSSGEGEASERGRSR-NH_2_; peptide 2: Ac-AARGSSDHAPYRRQGC-NH_2_) coupled to keyhole limpet hemocyanin (Eurogentec). An affinity purification column was generated by adding 0.96 mg of purified peptide 1 dissolved in coupling buffer (250 mM Tris pH 8.5, 25 mM EDTA) to 0.4 mL of SulfoLink resin (ThermoFisher) equilibrated in the same buffer. The resin was incubated with the peptide for two hours at 20°C with regular mixing, washed with 1.2 mL of coupling buffer and then blocked using 50 mM cysteine in coupling buffer at 20°C for 90 minutes with regular mixing. The resin was subsequently washed twice with 1 mL of 1 M NaCl, followed by another two washes with 5 mL of PBS. The immune serum was mixed with an equal volume of PBS and incubated with the peptide-coupled resin for 20 h at 4°C. The affinity-purified antibody was eluted in fractions using 100 mM glycine pH 2.5 into tubes containing 10× neutralization buffer (1M Tris pH 8.5, 2 M NaCl). Specificity of the antibody for use in immunoblots was tested by probing against cell lysates where pUL56 was absent or overexpressed, against lysates of cells infected with HSV-1 WT or ΔUL56, and against the GST-tagged purified recombinant protein (see below). BSA was added to the antibody for stabilization (final concentration 1 mg/mL) and the antibody was stored as a 50% (v/v) glycerol stock at −20°C.

#### Infection

Cell monolayers were infected with HSV-1 at the specified MOI diluted in complete media. For experiments to be analyzed by mass spectrometry, gradient-purified virus stocks were used. Otherwise, the infection was performed with crude virus stocks generated as described above. After adsorption for 1 h at 37°C with 5% CO_2_ and rocking every 15 min, the appropriate media was added to the well and this was designated 0 hpi. Infected cells were incubated at 37°C in a humidified 5% CO_2_ atmosphere until harvest.

#### Whole cell lysate preparation and protein digestion for quantitative temporal viromics

HaCaT cells were seeded into 6-well plates and infected in parallel at the specified MOI with gradient purified virus. At each indicated time point, cells were washed twice with PBS, and 250 µL lysis buffer was added (6M guanidine, 50 mM HEPES pH 8.5). Cell lifters (Corning) were used to scrape cells in lysis buffer, which was removed to an microcentrifuge tube, vortexed extensively, and then sonicated, and snap frozen in liquid nitrogen. After harvest, samples were stored at −70°C until all time points were harvested. Samples were thawed and cell debris was removed by centrifuging at 21,000×g for 10 min twice. Dithiothreitol (DTT) was added to a final concentration of 5 mM and samples were incubated for 20 min. Cysteines were alkylated with 14 mM iodoacetamide and incubated 20 min at room temperature in the dark. Excess iodoacetamide was quenched with DTT for 15 mins. Samples were diluted with 200 mM HEPES pH 8.5 to 1.5 M guanidine, followed by digestion at room temperature for 3 h with LysC protease (Wako) at a 1:100 protease-to-protein ratio. Samples were further diluted with 200 mM HEPES pH 8.5 to 0.5 M guanidine. Trypsin (Pierce) was then added at a 1:100 protease-to-protein ratio followed by overnight incubation at 37°C. The reaction was quenched with 5% (v/v) formic acid (FA), then centrifuged at 21,000×g for 10 min to remove undigested protein. Peptides were subjected to C18 solid-phase extraction (SPE; Sep-Pak, Waters) and vacuum-centrifuged to near-dryness.

#### Peptide labelling with tandem mass tags for whole cell experiments

In preparation for TMT labelling, desalted peptides were dissolved in 200 mM HEPES pH 8.5. Peptide concentration was measured by microBCA (Pierce), and >25 µg of peptide were labelled with TMT reagent. TMT reagents (0.8 mg) were dissolved in 43 μL anhydrous acetonitrile (MeCN) and 5 μL was added to the peptides at a final MeCN concentration of 30% (v/v). Sample labelling was as indicated in Table S1 and S4. Following incubation at room temperature for 1 h, the reaction was quenched with hydroxylamine to a final concentration of 0.5%. TMT-labelled samples were combined at a 1:1:1:1:1:1:1:1:1:1 ratio. The sample was vacuum-centrifuged to near dryness and subjected to C18 SPE (Sep-Pak, Waters). An unfractionated sample was analyzed initially to ensure similar peptide loading across each TMT channel, to avoid the need for excessive (>2-fold) electronic normalization. Samples were combined according to the correction factors from the unfractionated analysis and subjected to C18 SPE (Sep-Pak, Waters) and vacuum-centrifuged to near-dryness. The dried pellet was resuspended in 200 mM ammonium formate pH 10 and subjected to high pH reversed-phase (HpRP) fractionation is as described below.

#### Offline HpRP fractionation for whole cell lysate experiments

TMT-labelled tryptic peptides were subjected to HpRP fractionation using an Ultimate 3000 RSLC UHPLC system (Thermo Fisher Scientific) equipped with a 2.1 mm internal diameter (ID) x 25 cm long, 1.7 µm particle Kinetix Evo C18 column (Phenomenex). Mobile phase consisted of A: 3% (v/v) MeCN, B: MeCN and C: 200 mM ammonium formate pH 10. Isocratic conditions were 90% A / 10% C, and C was maintained at 10% throughout the gradient elution. Separations were conducted at 45°C. Samples were loaded at 200 µL/min for 5 min. The flow rate was then increased to 400 µL/min over 5 min, after which the gradient elution proceed as follows: 0-19% B over 10 min, 19-34% B over 14.25 min, 34-50% B over 8.75 min, followed by a 10 min wash at 90% B. UV absorbance was monitored at 280 nm and 15 s fractions were collected into 96-well microplates using the integrated fraction collector. Fractions were recombined orthogonally in a checkerboard fashion, combining alternate wells from each column of the plate into a single fraction, and commencing combination of adjacent fractions in alternating rows. Wells prior to the start or after the stop of elution of peptide-rich fractions, as identified from the UV trace, were excluded. This yielded two sets of 12 combined fractions, A and B, which were dried in a vacuum centrifuge and resuspended in 10 µL MS solvent (4% (v/v) MeCN / 5% (v/v) FA) prior to LC-MS3. For the time course experiment (Figure 1A) and ΔUL56/wildtype HSV-1 whole cell lysate experiment (Figure 5D), 12 set ‘A’ fractions were used for MS analysis.

#### Offline Tip-Based Strong Cation Exchange SCX Fractionation

Our previously described protocol for solid-phase extraction-based SCX peptide fractionation was modified for small peptide amounts (Dephoure and Gygi, 2011). Briefly, 10 mg of PolySulfethyl A bulk material (Nest Group Inc.) was loaded on to a fritted 200 µL tip in 100% Methanol using a vacuum manifold. SCX material was conditioned slowly with 1 mL SCX buffer A (7M KH_2_PO_4_, pH 2.65, 30% (v/v) MeCN), then 0.5 mL SCX buffer B (7 mM KH_2_PO_4_, pH 2.65, 350 mM KCl, 30% (v/v) MeCN) then 2 mL SCX buffer A. Dried peptides were resuspended in 500 mL SCX buffer A and added to the tip at a flow rate of ∼150 mL/min, followed by a 150 mL wash with SCX buffer A. Fractions were eluted in 150 µL buffer at increasing K^+^ concentrations (10, 25, 40, 60, 90, 150 mM KCl), vacuum-centrifuged to near dryness, then desalted using StageTips and vacuum-centrifuged to complete dryness and resuspended in 10 µL MS solvent (4% (v/v) MeCN / 5% (v/v) FA) prior to LC-MS3.

#### LC-MS/MS/MS for whole cell lysate experiments

Mass spectrometry data was acquired using an Orbitrap Lumos (Thermo Fisher Scientific, San Jose, CA). An Ultimate 3000 RSLC nano UHPLC equipped with a 300 µm ID x 5 mm Acclaim PepMap µ-Precolumn (Thermo Fisher Scientific) and a 75 µm ID x 50 cm 2.1 µm particle Acclaim PepMap RSLC analytical column was used. Loading solvent was 0.1% FA, analytical solvent A: 0.1% FA and B: 80% (v/v) MeCN + 0.1% FA. All separations were carried out at 55°C. Samples were loaded at 5 µL/min for 5 min in loading solvent before beginning the analytical gradient. The following gradient was used: 3-7% B over 3 min, 7-37% B over 173 min, followed by a 4-min wash at 95% B and equilibration at 3% B for 15 min. Each analysis used a MultiNotch MS3-based TMT method (McAlister et al., 2012; McAlister et al., 2014). The following settings were used: MS1: 380-1500 Th, 120,000 Resolution, 2×10^5^ automatic gain control (AGC) target, 50 ms maximum injection time. MS2: Quadrupole isolation at an isolation width of m/z 0.7, CID fragmentation (normalized collision energy (NCE) 35) with ion trap scanning in turbo mode from m/z 120, 1.5×10^4^ AGC target, 120 ms maximum injection time. MS3: In Synchronous Precursor Selection mode the top 6 MS2 ions were selected for HCD fragmentation (NCE 65) and scanned in the Orbitrap at 60,000 resolution with an AGC target of 1×10^5^ and a maximum accumulation time of 150 ms. Ions were not accumulated for all parallelizable time. The entire MS/MS/MS cycle had a target time of 3 s. Dynamic exclusion was set to +/-10 ppm for 70 s. MS2 fragmentation was trigged on precursors 5×10^3^ counts and above.

#### TMT Data analysis

In the following description, we list the first report in the literature for each relevant algorithm. Mass spectra were processed using a Sequest-based software pipeline for quantitative proteomics, “MassPike”, through a collaborative arrangement with Professor Steve Gygi’s laboratory at Harvard Medical School. MS spectra were converted to mzxml using an extractor built upon Thermo Fisher’s RAW File Reader library (version 4.0.26). In this extractor, the standard mzxml format has been augmented with additional custom fields that are specific to ion trap and Orbitrap mass spectrometry and essential for TMT quantitation. These additional fields include ion injection times for each scan, Fourier Transform-derived baseline and noise values calculated for every Orbitrap scan, isolation widths for each scan type, scan event numbers, and elapsed scan times. This software is a component of the MassPike software platform and is licensed by Harvard Medical School.

A combined database was constructed from (a) the human UniProt database (26th January, 2017), (b) HSV-1 strain KOS (Genbank entry JQ673480.1, manually updated with a single amino acid polymorphism in the ICP4 sequence identified in the KOS BAC strain used for virus generation), (c) common contaminants such as porcine trypsin and endoproteinase LysC. The combined database was concatenated with a reverse database composed of all protein sequences in reversed order. Searches were performed using a 20 ppm precursor ion tolerance (Haas et al., 2006). Product ion tolerance was set to 0.03 Th. TMT tags on lysine residues and peptide N termini (229.162932 Da) and carbamidomethylation of cysteine residues (57.02146 Da) were set as static modifications, while oxidation of methionine residues (15.99492 Da) was set as a variable modification.

To control the fraction of erroneous protein identifications, a target-decoy strategy was employed (Elias and Gygi, 2010). Peptide spectral matches (PSMs) were filtered to an initial peptide-level false discovery rate (FDR) of 1% with subsequent filtering to attain a final protein-level FDR of 1% (Kim et al., 2011; Wu et al., 2011). PSM filtering was performed using a linear discriminant analysis (Huttlin et al., 2010). This distinguishes correct from incorrect peptide IDs in a manner analogous to the widely used Percolator algorithm (Kall et al., 2007), though employing a distinct machine learning algorithm. The following parameters were considered: XCorr, ΔCn, missed cleavages, peptide length, charge state, and precursor mass accuracy. Protein assembly was guided by principles of parsimony to produce the smallest set of proteins necessary to account for all observed peptides (Huttlin et al., 2010). Where all PSMs from a given HSV-1 protein could be explained either by a canonical gene or non-canonical ORF, the canonical gene was picked in preference.

In three cases, PSMs assigned to a non-canonical ORFs or novel 6FT-ORFs were a mixture of peptides from the canonical protein and the ORF. This most commonly occurred where the ORF was a 5’-terminal extension of the canonical protein (thus meaning that the smallest set of proteins necessary to account for all observed peptides included the ORFs alone). In these cases, the peptides corresponding to the canonical protein were separated from those unique to the ORF, generating two separate entries.

Proteins were quantified by summing TMT reporter ion counts across all matching peptide-spectral matches using “MassPike”, as described (McAlister et al., 2012; McAlister et al., 2014). Briefly, a 0.003 Th window around the theoretical m/z of each reporter ion (126, 127n, 127c, 128n, 128c, 129n, 129c, 130n, 130c, 131n, 131c) was scanned for ions, and the maximum intensity nearest to the theoretical m/z was used. The primary determinant of quantitation quality is the number of TMT reporter ions detected in each MS3 spectrum, which is directly proportional to the signal-to-noise (S:N) ratio observed for each ion (Makarov and Denisov, 2009). Conservatively, every individual peptide used for quantitation was required to contribute sufficient TMT reporter ions (minimum of ∼1250 per spectrum) so that each on its own could be expected to provide a representative picture of relative protein abundance (McAlister et al., 2012). Additionally, an isolation specificity filter was employed to minimize peptide co-isolation (Ting et al., 2011). Peptide-spectral matches with poor quality MS3 spectra (more than 9 TMT channels missing and/or a combined S:N ratio of less than 250 across all TMT reporter ions) or no MS3 spectra at all were excluded from quantitation. Peptides meeting the stated criteria for reliable quantitation were then summed by parent protein, in effect weighting the contributions of individual peptides to the total protein signal based on their individual TMT reporter ion yields. Protein quantitation values were exported for further analysis in Excel (Microsoft).

For protein quantitation, reverse and contaminant proteins were removed, then each reporter ion channel was summed across all quantified proteins and normalized assuming equal protein loading across all channels. For further analysis and display in figures, fractional TMT signals were used (i.e. reporting the fraction of maximal signal observed for each protein in each TMT channel, rather than the absolute normalized signal intensity). This effectively corrected for differences in the numbers of peptides observed per protein. For TMT experiments, normalized S:N values are presented in Table S1 and S4 (‘Data’ worksheet).

Significance B was used to estimate the probability that each ratio was significantly different to 1 (Cox and Mann, 2008). Values were calculated and corrected for multiple hypothesis testing using the method of Benjamini-Hochberg in Perseus version 1.5.1.6 (Cox and Mann, 2008). A corrected p-value <0.05 was considered statistically significant. Hierarchical centroid clustering based on uncentered Pearson correlation of data normalized by comparing the signal:noise values to the average mock-infection were performed using Cluster 3.0 (Stanford University) and visualised using Java Treeview (http://jtreeview.sourceforge.net). For analysis of temporal classes, viral protein expression was normalised and subjected to K-means analysis using XLSTAT base (Addinsoft, version 18.06) and clustered with 1-15 classes.

#### Immunoblot of cell lysates

Cells were seeded into 24-well plates and infected with crude virus stocks in complete media or treated with 1 μg/mL doxycycline in complete media. Cells were harvested at the specified time point by scraping into the media and centrifuging at 16,000×g for 1 min. The cell pellet was resuspended in SDS loading buffer (50 mM Tris pH 6.8, 100 mM β-mercaptoethanol, 2% SDS, 10% glycerol). Samples were immediately boiled in a water bath for 5 min. Lysate from 1×10^5^ cells was used for sodium dodecyl sulfate-polyacrylamide gel electrophoresis (SDS-PAGE). Proteins were wet transferred onto 0.45 μm nitrocellulose membrane. After incubation with a primary antibody, secondary antibodies conjugated to an IRDye were used, and blots were visualized with an Odyssey CLx Imaging System (LiCor) using control software Image Studio v5.2.

#### Pathway analysis

The Database for Annotation, Visualisation and Integrated Discovery (DAVID) version 6.8 was used to determine pathway enrichment (Huang da et al., 2009). Proteins downregulated >2-fold were searched against a background of all proteins quantified, using default settings.

#### Immunoprecipitation

Monolayers of HEK 293T cells grown in 9 cm dishes (5×10^6^ cells/dish) were transfected using lipofectamine 2000 (Invitrogen) with GFP fusion proteins or GFP alone. For experiments with SILAC-labelled cells, the relevant labelled medium was used to prepare the transfection reagent. Cells were harvested 16-24 h post-transfection by scraping into the medium, pelleted (220×g, 5 min, 4°C) and washed three times with cold PBS. Cells were then lysed at 4°C in 1 mL lysis buffer (10 mM Tris pH 7.5, 150 mM NaCl, mM MgCl_2_, 0.5% Triton X-100, 1:100 Sigma protease inhibitors, 50 U/mL Sigma benzonase) for 45-90 min. The cell lysate was clarified by centrifugation (20,000×g, 10 min, 4°C), the supernatant transferred to fresh tubes, a BCA assay (Pierce) was performed to measure total protein concentration of clarified cell lysates, and samples were normalized (*input*).

GFP-Trap A beads (ChromoTek, 20 µL per sample) were washed three times by dilution in 800 µL wash buffer (10 mM Tris pH 7.5, 150 mM NaCl, 2 mM MgCl_2_, 0.05% Triton X-100), centrifugation (2500×g, 2 min, 4°C) to collect the beads and removal of the supernatant. Washed beads were incubated with the cleared lysate at 4°C on a rotating wheel for 45-70 min. The beads were collected by centrifugation and the supernatant (*unbound*) was removed. The beads were washed twice with 1 mL wash buffer, the supernatant was discarded, 45 µL of 2×SDS-PAGE loading buffer was added per experiment and the was mixture boiled at 95°C for 10 min to elute bound proteins. Samples were centrifuged again to sediment the beads (20,000×g, 2 min) and the supernatant (*bound*) was transferred to a fresh tube. Input, unbound and bound samples were separated by SDS-PAGE and analyzed by immunoblot. For mass spectroscopy analysis of SILAC samples, 8 µL of light-, medium-and heavy-labelled bound samples were mixed in a 1:1:1 ratio and frozen at −80°C until mass spectroscopy analysis.

#### Mass spectrometry of SILAC IP samples

Mass spectrometry analysis was performed by the proteomics facility of the University of Bristol (UK). Three biological repeats of each triple-labelled SILAC IP experiment were analyzed. Samples were run into precast SDS-PAGE gels for 5 minutes, the entire sample extracted from the gel as a single band, and then in-gel digested, reduced and alkylated using a ProGest automated digestion unit (Digilab). The resulting peptides were fractionated using an Ultimate 3000 nano-LC system in line with an Orbitrap Fusion Tribrid mass spectrometer (Thermo Scientific). In brief, peptides in 1% (v/v) FA were injected onto an Acclaim PepMap C18 nano-trap column (Thermo Scientific). After washing with 0.5% MeCN in 0.1% FA, peptides were resolved on a 250 mm × 75 μm Acclaim PepMap C18 reverse phase analytical column (Thermo Scientific) over a 150 min organic gradient using 7 gradient segments (1-6% solvent B over 1 min, 6-15% B over 58 min, 15-32% B over 58 min, 32-40% B over 5 min, 40-90% B over 1 min, held at 90% B for 6 min and then reduced to 1% B over 1min) with a flow rate of 300 nL per minute. Solvent A was 0.1% FA and solvent B was aqueous 80% MeCN in 0.1% FA. Peptides were ionized by nano-electrospray ionization at 2.0 kV using a stainless steel emitter with an internal diameter of 30 µm (Thermo Scientific) and a capillary temperature of 275°C. All spectra were acquired using an Orbitrap Fusion Tribrid mass spectrometer controlled by Xcalibur 2.1 software (Thermo Scientific) and operated in data-dependent acquisition mode. FTMS1 spectra were collected at a resolution of 120,000 over a scan range (m/z) of 350-1550, with an automatic gain control (AGC) target of 300,000 and a max injection time of 100 ms. Precursors were filtered using an Intensity Range of 1×10^4^ to 1×10^20^ and according to charge state (to include charge states 2-6) and with monoisotopic precursor selection. Previously interrogated precursors were excluded using a dynamic window (40 s +/-10 ppm). The MS2 precursors were isolated with a quadrupole mass filter set to a width of 1.4 m/z. ITMS2 spectra were collected with an AGC target of 20,000, max injection time of 40 ms and CID collision energy of 35%.

The raw data files were processed using MaxQuant v. 1.5.7.4 (Cox and Mann, 2008). The in-built Andromeda search engine (Cox et al., 2011) was used to search against the human and HSV-1 strain KOS proteomes as used for TMT analysis (above). Trypsin/P digestion, standard modifications (oxidation, N-terminal acetylation) were selected as group-specific parameters and SILAC quantification was performed using light (Arg0, Lys0), medium (Arg6, Lys4) and heavy (Arg10, Lys8) labels. Re-quantification, razor protein FDR, and second peptide options were enabled for the processing. The quantified data were analyzed with Perseus v. 1.6.1.2 (Tyanova et al., 2016) using the normalized ratios obtained by MaxQuant. Proteins only identified by site or against the reverse database, as well as common experimental contaminants such as keratins (specified in the MaxQuant contaminants file), were removed and the experiments grouped by biological repeat. Only proteins identified in at least two of the three biological repeats were considered for analysis. A one-sample, two-sided t-test with a threshold p-value of 0.05 was performed on each group to identify significantly enriched proteins. Proteins with a log_2_ fold change greater than 1 and a p value smaller than 0.05 were designated as potential interactors of pUL56.

#### Recombinant protein expression and purification

For bacterial recombinant expression, the cytoplasmic region (residues 1-207) of UL56 from HSV-1 strain KOS was cloned into a vector derived from pOPT (Teo et al., 2004) encoding *Schistosoma japonicum* GST followed by a human rhinovirus 3C cleavage sequence fused to the N terminus and LysHis_6_ fused to the C terminus (GST-UL56(1-207)-His). Full-length (residues 1-454) and truncated forms (residues 1-362, 27-362, 27-275, 276-362 and 27-236) of GOPC (UniProt ID Q9HD26-2) were cloned from HeLa cell cDNA into a vector derived from pOPT (Teo et al., 2004) encoding a MetAlaHis_6_ tag fused to the N terminus of each construct (His-GOPC).

His-GOPC (both full-length and truncations) was expressed in *Escherichia coli* BL21(DE3)pLysS cells (Novagen) and GST-UL56(1–207)-His was expressed in *E. coli* T7 Express LysY/Iq cells (New England Biolabs). Cells were cultured in 2×TY medium to an OD600 between 0.8 and 1.0. For His-GOPC, the culture was cooled to 22°C before adding 0.2 mM IPTG and culturing for a further 16 h. For GST-UL56(1–207)-His, 1 mM IPTG was added and the cells were cultured for a further 2 h. Cells were harvested by centrifugation and pellets stored at −80°C.

For His-GOPC, cell pellets were resuspended on ice in Ni^2+^ wash buffer (20 mM Tris pH 7.5, 20 mM Imidazole, 500mM NaCl) supplemented with 0.5 mM MgCl_2_, 1.4 mM 2-mercaptoethanol, 0.05% TWEEN-20, 400 U Bovine DNAse I and 200 µL EDTA-free protease inhibitors (Sigma-Aldrich) and lysed by passing through a TS series cells disruptor (Constant Systems) at 24 kpsi. Lysates were cleared by centrifugation (40,000×g, 30 min, 4°C) and incubated with NiNTA agarose (Qiagen) pre-equilibrated in Ni^2+^ wash buffer for 60 min at 4°C. The resin was washed with >20 column volumes (cv) of Ni^2+^ wash buffer and protein was eluted in Ni^2+^ elution buffer (20 mM Tris pH 7.5, 250 mM imidazole, 500mM NaCl) before being concentrated and applied to a Superdex 200 16/600 gel filtration column that had been pre-equilibrated in gel filtration buffer (20 mM Tris, 200 mM NaCl, 1 mM DTT) at room temperature. Eluted fractions containing purified His-GOPC were pooled, concentrated and small (<100 uL) aliquots were snap-frozen in liquid nitrogen for storage at −80°C.

For GST-UL56(1-207)-His, cells were resuspended on ice in 50 mM sodium phosphate pH 7.6, 300 mM NaCl, 0.5 mM MgCl_2_, 1.4 mM 2-mercaptoethanol, 0.05% TWEEN-20, 400 U Bovine DNAse I and 200 µL EDTA-free protease inhibitors (Sigma-Aldrich) before lysis and clarification as described above. Cleared lysates were incubated with glutathione Sepharose 4B (GE Life Science) that had been pre-equilibrated in GSH wash buffer (50 mM sodium phosphate pH 7.6, 300 mM NaCl, 1 mM DTT) for 1 h at 4°C. The resin was washed with 10 cv of GSH wash buffer before being resuspended in 20 cv of 25 mM sodium phosphate pH 7.5, 150 mM NaCl, 1 MgCl_2_, 0.5 mM DTT and incubated at room temperature for 30 min with 50 U/mL benzonase nuclease (Sigma-Aldrich) to digest co-purifying nucleic acids. The resin was then washed with 20 cv of 50 mM sodium phosphate pH 7.6, 1 M NaCl to remove residual nucleotide binding before being washed with a further 40 cv of GSH wash buffer. Protein was eluted using GSH wash buffer supplemented with 25 mM reduced glutathione. The protein was then captured using NiNTA agarose that had been equilibrated in Ni^2+^ wash buffer, the resin was washed with >20 cv of Ni^2+^ wash buffer, and the protein eluted in Ni^2+^ elution buffer before being injected onto a 10/300 Superdex 200 gel filtration column (GE Healthcare) equilibrated in gel filtration buffer (as above). Eluted fractions containing UL56 were pooled, concentrated and snap-frozen in small (<100 µL) aliquots for storage at −80°C.

#### Protein GST pull-down assays

Bait proteins were diluted to 5 µM in pull-down buffer (20 mM Tris pH 7.5, 200 mM NaCl, 0.1% NP-40, 1 mM DTT, 1 mM EDTA) and, for each experiment, 200 µL of bait mixture was incubated for 15-30 min at room temperature with 10 µL of glutathione magnetic beads (Pierce) that had been pre-equilibrated in pull-down buffer. Supernatant was removed and resin was washed twice with pull-down buffer. Bait-loaded resin was incubated with purified His-GOPC (full-length or truncated) or clathrin N-terminal domain (Muenzner et al., 2017) diluted to 10 µM in pull-down buffer for 60 min at room temperature in a final volume of 200 µL per experiment. Unbound prey was removed and the beads washed four times with pull-down buffer. Bound proteins were eluted using pull-down buffer supplemented with 50 mM reduced glutathione. Samples were resolved by SDS-PAGE and visualized using InstantBlue Coomassie stain (Expedeon).

#### Immunofluorescence microscopy

Cells were seeded to be a third confluent on #1.5 glass coverslips and transfected with TransIT-LT1 or infected at MOI of 1 with crude virus stocks in complete media. At 1 day post-transfection or 6 hpi, the samples were fixed in 3% (v/v) formaldehyde PBS for 15 min at room temperature. Cells were permeabilized and washed using PBS supplemented with 1% (v/v) FBS, 0.1% Triton-X100. If the primary antibody was from a rabbit, a 2 h blocking step using PBS supplemented with 100 μg/mL human IgG, 10% (v/v) FBS was included before incubation with the primary antibody. Antibodies were diluted into PBS supplemented with 10% (v/v) FBS (plus 100 μg/mL human IgG for antibodies raised in rabbit). After immunostaining, the coverslips were mounted with ProLong Gold Antifade Mountant containing 4’,6-diamidino-2-phenylindole (DAPI) (ThermoFisher). Samples were analyzed with an inverted Olympus IX81 widefield microscope. Illumination was performed with a Lumen 200 arc lamp (Prior Scientific) and bandpass filters for DAPI (excitation of 350/50 nm and emission of 455/50 nm), Alexa Fluor 488 (excitation of 490/20 nm and emission of 525/36 nm), and Alexa Fluor 568 (excitation of 572/35 nm and emission of 605/52 nm) (Chroma Technology Corp). Images were acquired with Image-Pro Plus software (Media Cybernetics), a Retiga EXi Fast1394 interline CCD camera (QImaging), and a 60x PlanApo N oil objective (numerical aperture, 1.42) (Olympus) for a pixel resolution of 107.5 nm/pixel.

#### Generation of Flp-In™ T-REx™-293 stable cells

The pUL56-inducible cell line was generated according to the manufacturer’s instructions. Briefly, pUL56 was cloned into pcDNA5/FRT/TO. Flp-In™ T-REx™-293 cells were transfected with pcDNA5/FRT/TO-pUL56 and the Flp recombinase expression plasmid pOG44 using TransIT-LT1. One day post-transfection, the cells were selected in 100 μg/mL hygromycin and 15 mg/mL blasticidin. Single cell clones were then isolated and screened for pUL56 expression. Expression of pUL56 was induced by incubating cells with 1 μg/mL doxycycline 1 day prior to harvest.

#### Virus growth curves, and plaque assays

Growth curves were performed using HaCaT cells infected in complete media with crude virus stocks of HSV-1 WT or HSV-1 ΔUL56 at MOI of 10. After adsorption for 1 h at 37°C, cells were incubated with acid wash (40 mM citric acid, 135 mM NaCl, 10 mM KCl; pH 3.0) for 1 min and washed 3x with PBS before cell culture media was added back. The time of acid wash was deemed 0 hpi. At various times post-infection, cells were harvested by freezing the plate at −70°C. After freezing the last time point, samples were freeze-thawed together 2 subsequent times and scraped before they were titred. Titrations were performed on Vero monolayers. Cells were inoculated with serial dilutions of the samples for 1 h, after which DMEM containing 0.3% high viscosity carboxymethyl cellulose, 0.3% low viscosity carboxymethyl cellulose, 2% (v/v) FBS, 2 mM L-glutamine, 100 U/mL penicillin, and 100 μg/mL streptomycin was overlaid. After 3 days, cells were fixed in 3.75% (v/v) formaldehyde in PBS for 30 min and stained with 0.1% toluidine blue.

For plaque size measurements, HaCaT, HFF hTERT, or Vero cells were grown in 6-well plates. The cells were infected and fixed as described above, but they were stained with an anti-gD antibody (LP2). Plaques were visualized with a secondary antibody conjugated to horseradish peroxidase and the DAB peroxidase substrate following the manufacturer’s instructions (Vector SK4105). Plaques were scanned at 300 dpi and plaque diameters were measured with ImageJ (https://imagej.nih.gov/ij/).

#### Plasma membrane profiling

SILAC labelled HaCaT cells (as described above) were grown in 15 cm dishes and infected with gradient purified HSV-1 WT or HSV-1 ΔUL56 or mock infected in complete media at MOI 10. Plasma membrane profiling was performed as described previously with minor modifications (Weekes et al., 2010). Briefly, at 6 hpi cells were washed, surface sialic acid residues oxidized with sodium-meta-periodate and labelled with aminooxy-biotin. The reaction was quenched and the biotinylated cells scraped into 1% (v/v) Triton X-100 lysis buffer. Biotinylated glycoproteins were enriched with high affinity streptavidin agarose beads and washed extensively. Captured protein was denatured with DTT, alkylated with iodoacetamide (IAA) and digested on-bead with trypsin in 100 mM HEPES pH 8.5 for 3 h. Tryptic peptides were collected and fractionated by tip-based SCX strong cation exchange, generating six fractions for MS analysis.

#### LC-MS/MS for plasma membrane experiments

Mass spectrometry data was acquired using an Orbitrap Lumos (Thermo Fisher Scientific, San Jose, CA). An Ultimate 3000 RSLC nano UHPLC equipped with a 300 µm ID x 5 mm Acclaim PepMap µ-Precolumn (Thermo Fisher Scientific) and a 75 µm ID x 50 cm 2.1 µm particle Acclaim PepMap RSLC analytical column was used. Loading solvent was 0.1% FA, analytical solvent A: 0.1% FA and B: 80% (v/v) MeCN + 0.1% FA. All separations were carried out at 55°C. Samples were loaded at 5 µL/min for 5 min in loading solvent before beginning the analytical gradient. The following gradient was used: 3-7% B over 4 min, 7-37% B over 116 min, followed by a 4-min wash at 95% B and equilibration at 3% B for 15 min. Each analysis used an MS2 DDA acquisition using the following settings: MS1: 375-1500 Th, 60,000 Resolution, 4×10^5^ automatic gain control (AGC) target, 50 ms maximum injection time. MS2: Quadrupole isolation at an isolation width of m/z 1.6, HCD fragmentation (normalised collision energy (NCE) 35) with ion trap scanning in rapid mode from m/z 110, 1×10^4^ AGC target, 35 ms maximum injection time.

The resulting spectra were processed in Maxquant 1.5.8.3 using medium (Arg6, Lys4) and heavy (Arg10, Lys8) labels. Data was searched against the human and HSV-1 strain KOS proteomes as used for TMT analysis (above). Carbamidomethyl (C) was set as a fixed modification, oxidation (M) and acetylation (protein N termini) set as variable modifications. Protein and peptide FDR were both set to 0.01, re-quantify was enabled and minimum ratio count was set to 2. Hierarchical centroid clustering based on uncentered Pearson correlation of the normalised ratios generated by MaxQuant was performed using Cluster 3.0 (Stanford University) and visualised with Java Treeview (http://jtreeview.sourceforge.net).

#### Generation of CRISPR knockout HaCaT cells

HaCaT cells were seeded at 50% confluence and transfected with the PX459 CRISPR plasmid containing relevant guide RNAs (GOPC 1: GGAACATGGATACCCCGCCA; GOPC 2: GAGAGATCGATCCAGACCAAG) and Lipofectamine 2000 according to the manufacturer’s instructions. pSpCas9(BB)-2A-Puro (PX459) V2.0 was a gift from Feng Zhang (Addgene plasmid # 62988; http://n2t.net/addgene:62988; RRID:Addgene_62988) (Ran et al., 2013). One day post-transfection the medium was changed to contain 2 μg/mL puromycin, and 3 days post-transfection the medium was changed to selection-free medium. Clonal cell lines were expanded and tested for loss of GOPC by western blot analysis and genomic sequencing.

#### Flow Cytometry

HaCaT cells infected with crude virus stocks were washed 2 times with PBS and detached with accutase. Cells were pelleted at 400×g for 5 min and washed once with PBS containing 2% (v/v) FBS. For extracellular staining, cells were stained with anti-human CD282 (TLR2) antibody (BioLegend, 153003) and incubated for 1 h at room temperature. Stained cells were washed once and fixed in 4% (v/v) formaldehyde in PBS for 20 min at room temperature. Data was acquired with a FACSCalibur and analyzed with Flowing Software version 2.5.1 (http://flowingsoftware.btk.fi/).

#### RT PCR quantification

Cells infected with crude virus stocks were harvested by scraping and centrifugation at 200×g for 5 min. They were washed once in PBS before lysis buffer was added (4 M guanidinium thiocyanate, 25 mM Tris pH 7). After 10 min on ice, an equal volume of 100% ethanol was added and this was loaded onto a spin column. Columns were washed once with 1 M guanidinium thiocyanate, 25 mM Tris pH 7, 10% (v/v) ethanol and twice with 25 mM Tris pH 7, 70% (v/v) ethanol before being eluted with water. Samples were treated with RQ1 DNase according to the manufacturer’s instructions with RNaseOUT. Reverse transcription was performed with M-MLV RT and a random hexamer primer mix, and PCR was carried out with Phusion DNA polymerase. RT PCR products were visualized with a 1% (w/v) agarose TAE gel with 1 μg/mL ethidium bromide. Image acquisition was achieved with a G:BOX gel imager with control software GeneSnap v7.12.06.

#### Data availability

The mass spectrometry proteomics data will be deposited with the ProteomeXchange Consortium (http://www.proteomexchange.org/) via the PRIDE partner repository under the data set identifier PXDxxxxxx (http://www.ebi.ac.uk/pride/archive/projects/PXDxxxxxx).

Unprocessed peptide data files for Figures 1, 3, 5 and 6 are available at doi: https://data.mendeley.com/datasets/g5sf93zwtf/1

