## Supplemental Figures S1-S4 for "Herpes simplex virus-1 pUL56 degrades GOPC to alter the plasma membrane proteome"

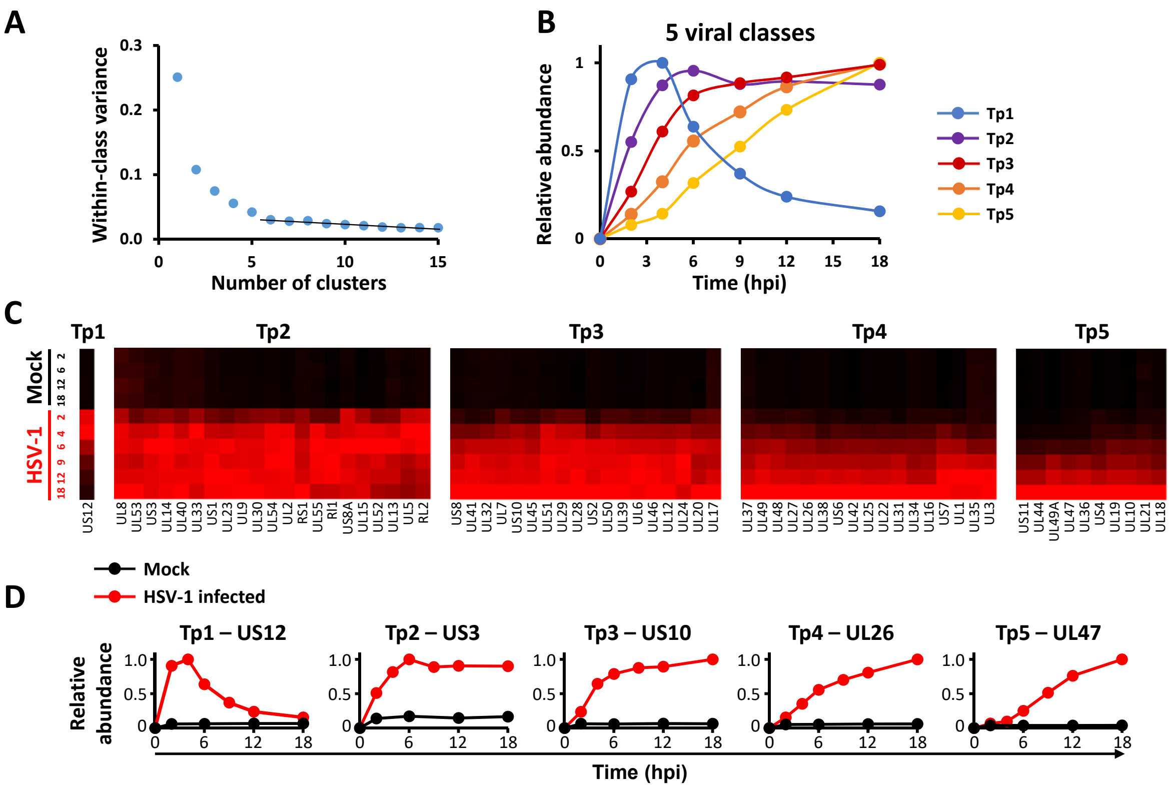


<u>Figure S1, related to Figure 1: Definition of temporal classes of HSV-1 gene expression</u>

**A**. Number of temporal classes of HSV-1 gene expression. The k-means approach was used with 1-15 classes to cluster viral proteins, and the summed distance of each protein from its cluster centroid was calculated. Although this summed distance necessarily becomes smaller as more clusters are added, the rate of decline decreases with each added group, eventually settling at a fairly constant rate of decline that reflects overfitting; clusters added prior to this point reflect underlying structure in the temporal protein data, whereas clusters subsequently added through overfitting are not informative. The point of inflexion fell between 5 to 6 classes, suggesting that there are at least 5 distinct temporal protein profiles of viral protein expression. **B**. Average temporal profiles for each class. **C**. Temporal profiles of proteins in each k-means class were subjected to hierarchical clustering by Euclidian distance. **D**. Temporal profiles of the central protein from each cluster.

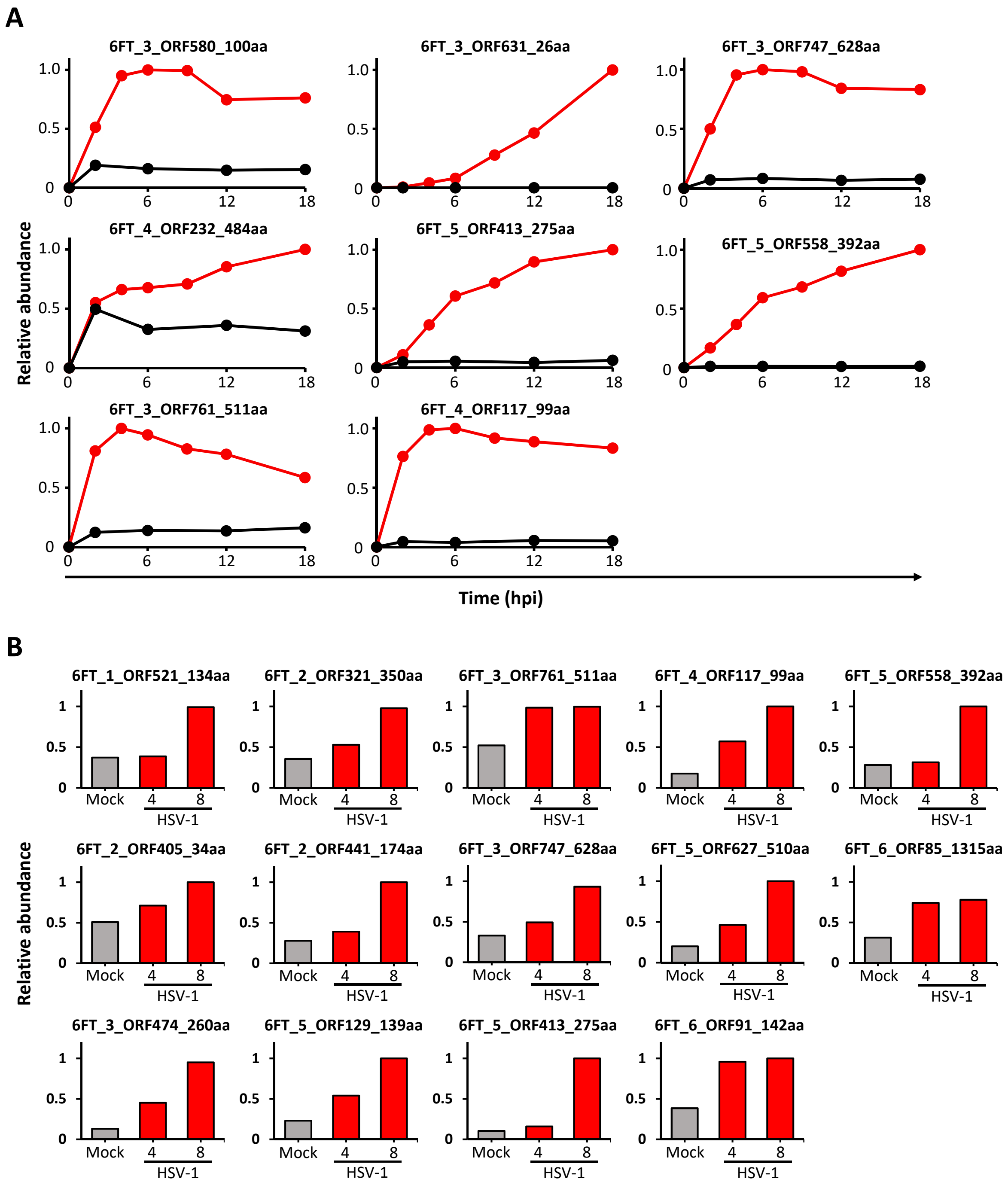


Figure S2, related to Figure 1 and Figure 5: Profiles of novel non-canonical ORFs identified by proteomics.

**A**. Temporal profiles of novel non-canonical ORFs identified in the HSV-1 time course (Figure 1). Proteomic data was analyzed against a database containing all ORFs of ≥8 amino acids from a six-frame translation of the HSV-1 sequence. The profiles shown are from identified peptides unique to a novel six frame translation sequence. Protein profiles that did not show a >1.5-fold increase in any HSV-1 infected samples were removed. **B**. Temporal profiles of novel non-canonical ORFs identified from the comparison of HSV-1 WT to HSV-1 ΔUL56 virus (Figure 5). The results shown are from all of the quantified non-canonical ORFs that increased in abundance over the course of infection. Protein profiles that did not show a >1.5-fold increase in any HSV-1 infected samples were removed.

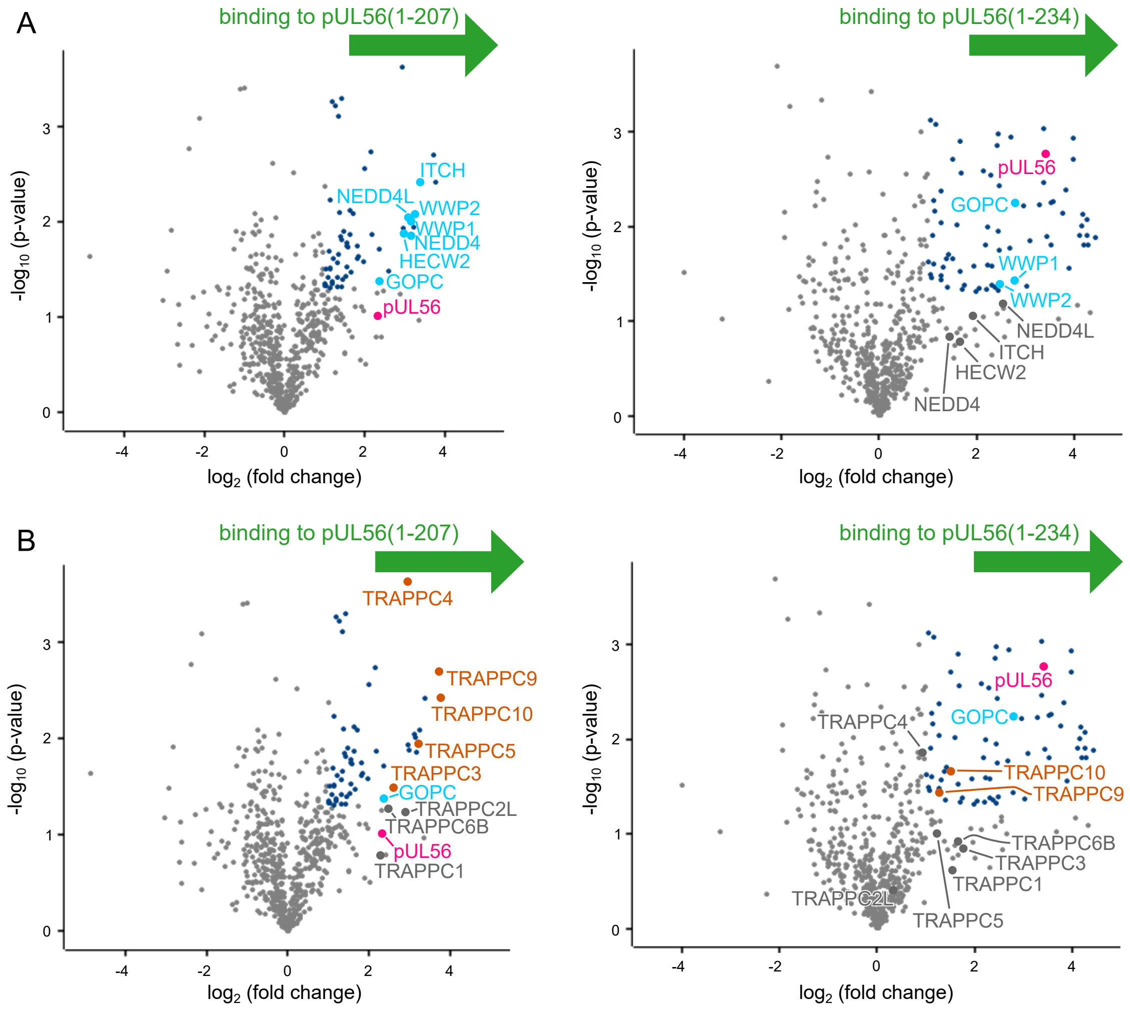


<u>Figure S3, related to Figure 3: The pUL56 cellular interactome identified by SILAC-based IP</u>

SILAC-labelled HEK293T cells were transfected with full-length pUL56 (residues 1-234) N-terminally tagged with GFP, or the cytoplasmic domain (residues 1-207) of pUL56 C-terminally tagged with GFP, or with GFP alone, and subjected to immunoprecipitation (IP) using a GFP affinity resin. The volcano plot shows average fold enrichment in IP of GFP-tagged pUL56 versus GFP against significance across three biological replicates. Significantly enriched proteins (>2-fold enrichment and $p < 0.05$) are colored blue and selected proteins are highlighted. **A**. Both pUL56 cytoplasmic domain (left, as in Figure 3) and the full-length protein (right) interact with GOPC and NEDD4 family ubiquitin ligases. **B**. GFP-tagged pUL56 binds members of the TRAPPII complex. TRAPPII complex specific components TRAPPC9 and TRAPPC10 were the proteins with most enhanced enrichment in the IP of GFP-tagged pUL56 cytoplasmic domain (left) and all members of the TRAPPII complex were identified as highly enriched, although three (TRAPPC6B, TRAPPC2L and TRAPPC1) were below the significance threshold. TRAPPC9 and TRAPPC10 are also significantly enriched in the IP of full-length pUL56 (right). The same dataset is depicted in (**A**) and (**B**).

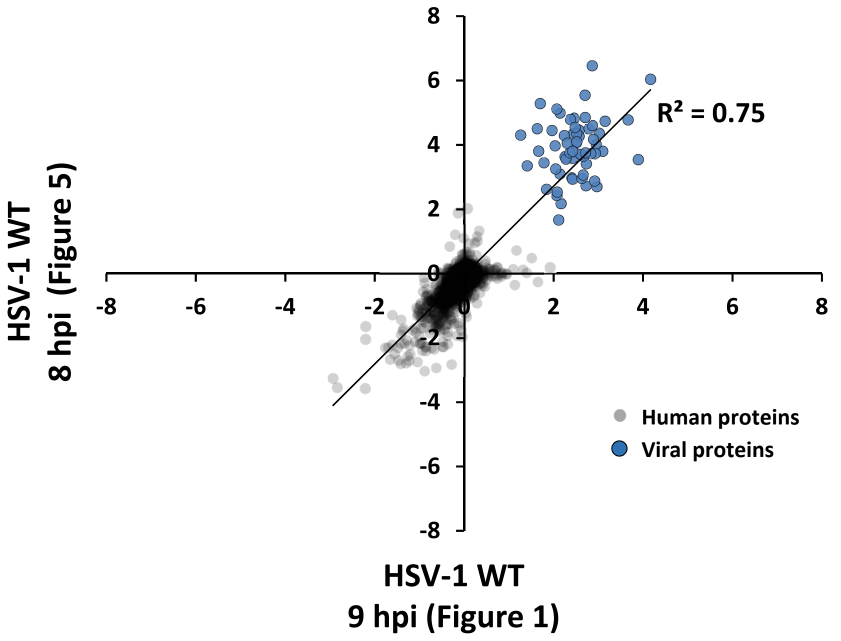


Figure S4, related to Figure 1 and Figure 5: Correlation of proteomic profiling between repeat experiments.

Scatter plot showing the correlation between repeat experiments: fold-change 9 hpi (Figure 1) and 8 hpi (Figure 5) of HSV-1 WT compared to mock.
